# Targeting Transferrin Receptor to Transport Antisense Oligonucleotides Across the Blood-Brain Barrier

**DOI:** 10.1101/2023.04.25.538145

**Authors:** Scarlett J Barker, Mai B Thayer, Chaeyoung Kim, David Tatarakis, Matthew Simon, Rebekah L Dial, Lizanne Nilewski, Robert C Wells, Yinhan Zhou, Megan Afetian, Alfred Chappell, Kylie S Chew, Johann Chow, Allisa Clemens, Claire B Discenza, Jason Dugas, Chrissa Dwyer, Timothy Earr, Connie Ha, David Huynh, Srini Jayaraman, Wanda Kwan, Cathal Mahon, Michelle Pizzo, Elysia Roche, Laura Sanders, Alexander Stergioulis, Raymond Tong, Hai Tran, Joy Zuchero, Anthony A Estrada, Kapil Gadkar, Christopher MM Koth, Pascal E Sanchez, Robert G Thorne, Ryan J Watts, Thomas Sandmann, Lesley Kane, Frank Rigo, Mark S Dennis, Joseph W Lewcock, Sarah L DeVos

## Abstract

Antisense oligonucleotides (ASO) are promising therapies for neurological disorders, though they are unable to cross the blood-brain barrier (BBB) and must be delivered directly to the central nervous system (CNS). Here, we use a human transferrin receptor (TfR)-binding molecule to transport ASO across the BBB in mice and non-human primates, termed oligonucleotide transport vehicle (OTV). Systemically delivered OTV drives significant, cumulative, and sustained knockdown of the ASO target across multiple CNS regions and all major cell types. Further, systemic OTV delivery enables more uniform ASO biodistribution and knockdown compared to two other clinically relevant ASO delivery routes: a standard, high affinity TfR antibody, or direct ASO delivery to the CSF. Together, our data support systemically delivered OTV as a potential therapeutic platform for neurological disorders.

**One-Sentence Summary:** Systemically dosed OTV delivered via TfR1 targeting shows widespread and cumulative target knockdown in the mouse and NHP CNS.

## Main Text

Oligonucleotide based therapeutics – both antisense oligonucleotides (ASO) and short interfering RNAs (siRNA) – have become increasingly used to target various neurological disorders based on their ability to selectively modulate targets that are difficult to influence with other modalities [1–4]. Following the approval of Nusinersen in 2016 for Spinal Muscular Atrophy (SMA) and encouraging data from other ASO targets [5], numerous oligonucleotide-based therapies targeting central nervous system (CNS) disorders have entered clinical trials[6, 7]. Due to their inherent biophysical properties (e.g., size, charge, backbone chemistry), oligonucleotides clear from circulation quickly and are unable to efficiently cross the blood-brain barrier (BBB) [8, 9], thus requiring repeat intrathecal (IT) or intracerebroventricular (ICV) dosing directly into the cerebrospinal fluid (CSF) to access the CNS [10, 11]. IT oligonucleotide delivery is not without its limitations, including diffusion-mediated biodistribution that results in reduced drug levels in deeper brain regions and increased levels in lumbar spinal cord [12], as well as the potential for adverse events related to IT dosing that can lead to dose level limitations or even study discontinuation[6, 13]. Additionally, the delivery of ASOs and siRNAs to most peripheral non-hepatic or renal tissues, such as muscle, are similarly limited due to inefficient functional uptake[7, 14].

Several technologies have emerged to enable less invasive routes of administration and/or facilitate delivery of therapeutics to the CNS with varying degrees of success, including cholesterol-conjugates, peptide-conjugates, aptamers, and sugars [15, 16]. High physiological expression of transferrin receptor 1 (TfR1) on brain endothelial cells at the BBB has led several groups to pursue CNS drug delivery via TfR1 mediated transcytosis [17–22]. Our previous work led to the development of the transport vehicle (TV), a platform in which low affinity human TfR1 (hTfR1) binding has been engineered into an Fc domain of the TV to deliver multiple types of cargo across the BBB, including antibodies[23, 24], enzymes[25–27], and other proteins[28]. Across modalities, enhanced brain uptake with uniform distribution throughout the CNS is a defining hallmark of the TV. Various TVs are currently in human clinical trials, highlighting the potential for clinical translatability of the TV platform. However, given that oligonucleotide-based therapeutics require access to a cell’s cytosol or nucleus, it is unclear whether TfR-mediated approaches like the TV would be capable of delivering an oligonucleotide therapeutic modality to the necessary site of action for target engagement. Here we generated a modular platform using an engineered TV conjugated to an ASO for delivery of therapeutic molecules to the brain, termed oligonucleotide transport vehicle (OTV). We demonstrate that OTV can successfully cross the BBB following systemic intravenous (IV) administration and provide ASO-mediated target RNA knockdown in the mouse and non-human primate (NHP) brain. Further, OTV differentiates from oligonucleotide delivery via conjugation to a standard bivalent TfR antibody as well as direct CSF administration, the current standard of care for CNS targeted oligonucleotides.

## Results

### Site Specific ASO Conjugation to the TV Platform

To examine the impact of hTfR1 binding on ASO biodistribution in the CNS and periphery, we employed a tool ASO designed against *Malat1* [29], a long non-coding RNA. *Malat1* is a common oligonucleotide target for platform development[30] and poses several benefits for testing with the OTV platform, including 1) high basal expression levels for ease of measuring appreciable knockdown, 2) easily detectable expression across all CNS cell types and peripheral tissues for organism-wide pharmacodynamic assessment[31], and 3) exclusively nuclear RNA localization[32] to determine whether the ASO portion of the OTV molecule can not only escape the endolysosomal system, but enable transport into the nucleus where ASOs enact target RNA knockdown through RNAase H1 recruitment [33]. The ASO used for all *in vitro* studies and *in vivo* mouse studies against murine *Malat1* is subsequently termed ASO.

The TV used for all OTV studies is a huIgG containing an engineered Fc domain that binds with high specificity to a single apical domain of hTfR1 while not impeding normal transferrin binding [23] (Fig. 1A). The Fab domain of OTV does not bind known antigens in rodents or NHP and was designed to be non-targeting in order to understand hTfR1 mediated biodistribution without binding Fabs impacting distribution. To control the oligonucleotide:TV ratio (OTR), we used a knob-in-hole Fc to introduce a single engineered cysteine at serine 239 of the Fc hole to enable conjugation through a maleimide-C6 linker attached at the 5’ end of the ASO (Fig. 1B) [34]. Following conjugation, a desired OTR of 1 was confirmed via mass spectrometry and size exclusion chromatography (SEC) (Fig. 1C; Fig. S1A). To test the impact of ASO conjugation on OTV binding to hTfR1, we first measured binding of OTV to hTfR1 in HEK cells. Comparison to a TV control (no ASO) revealed that the presence of the ASO does not impact cell binding (Fig. 1D). Using surface plasmon resonance (SPR) to measure hTfR1 affinity directly, we again observed no impact with OTV versus the TV control (Fig. 1E-F). These data highlight the feasibility of achieving site-specific ASO conjugation to generate OTV without impacting hTfR1 binding.

**Fig 1.**
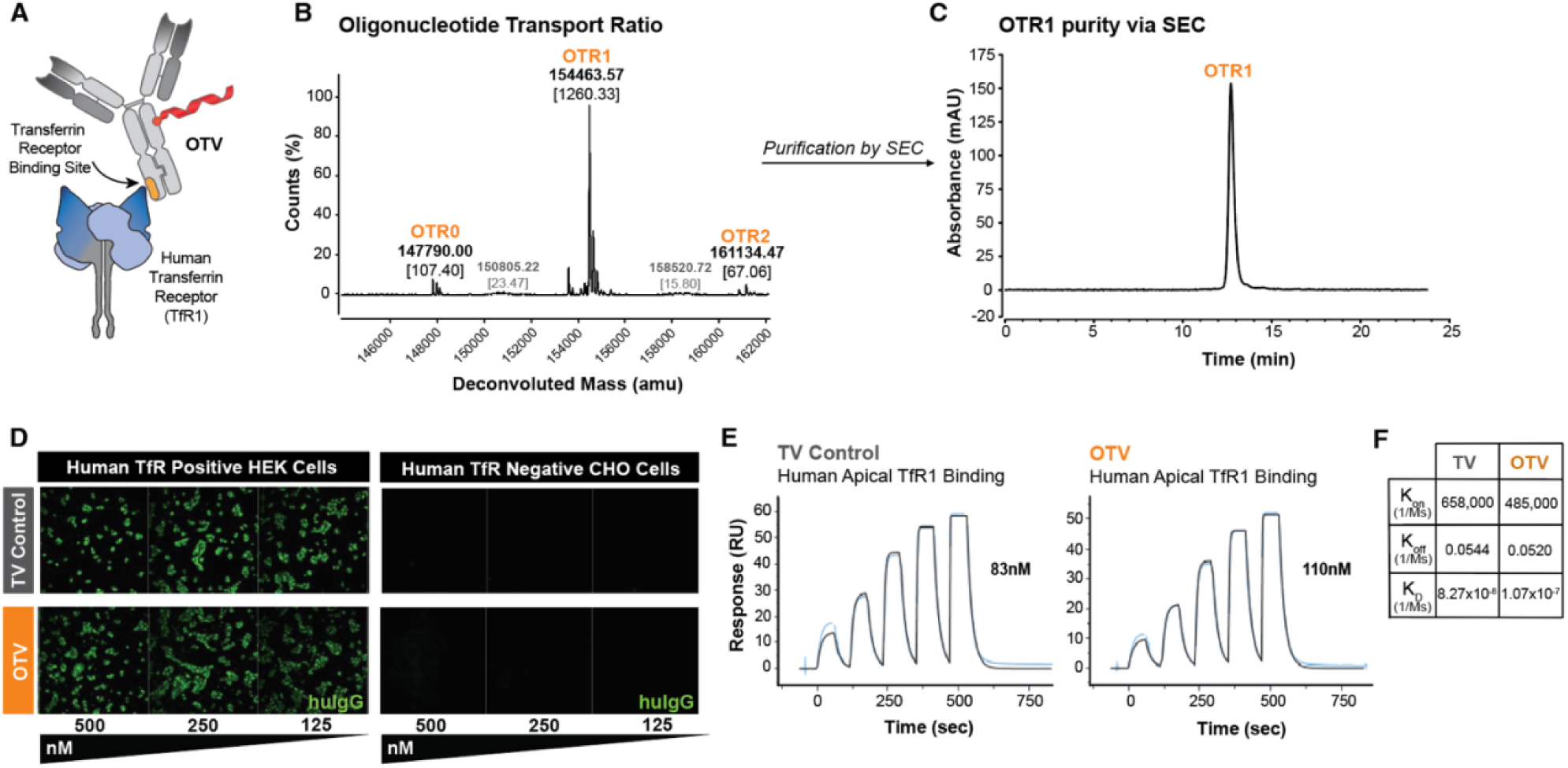
Site-specific ASO Conjugation to OTV does not alter binding to hTfR1. **(A)** Schematic of OTV binding to the apical domain of hTfR1. (**B)** Mass spectrometry shows successful ASO conjugation to OTV with a majority of OTV having the desired OTR1. (**C)** Following further purification, SEC data confirms purified OTR1. (**D)** No impact of OTV on successful hTfR1 binding compared to a non-ASO TV control demonstrated by an *in vitro* uptake assay in cells that express hTfR1 (HEK, left panels) and cells that do not (CHO, right panels). (**E-F)** Binding of the TV control (left) and OTV (right) to hTfR1 by SPR shows similar binding affinity (E) and dissociation constants (F).

### OTV Drives ASO Cellular Uptake in vitro through hTfR1 binding

To understand the impact of ASO conjugation on cellular trafficking, we conjugated a Cy3-labeled ASO to the TV huIgG, then labeled the TV with Alexa-647 to create a double-labeled OTV. Of note, the Cy3 addition to the ASO does not impede *Malat1* RNA knockdown (Fig. S1B). We then monitored cellular uptake and subcellular distribution of the double-labeled OTV as compared to naked ASO in SH-SY5Y cells, a commonly used cell line of neural origin. Both naked ASO and ASO derived from OTV were robustly internalized (Fig. 2A) and trafficked through the endolysosomal system with predominant localization in lysosomes after 24 hours (Fig. 2B-C; Fig. S1C). Cellular uptake and trafficking of the huIgG portion of OTV was not substantially altered by conjugation to ASO (Fig. 2D-F; Fig. S1D). Super resolution microscopy confirmed the localization of naked ASO and OTV-derived ASO in late endosomes (Rab7^+^) and lysosomes (Lamp1^+^) in SH-SY5Y cells (Fig. 2G). Following OTV treatment, both ASO and huIgG were found together in these compartments as shown by the overlap of ASO (purple) and huIgG (white) with markers of both endosomes and lysosomes (green) (Fig. 2G, right).

**Fig 2.**
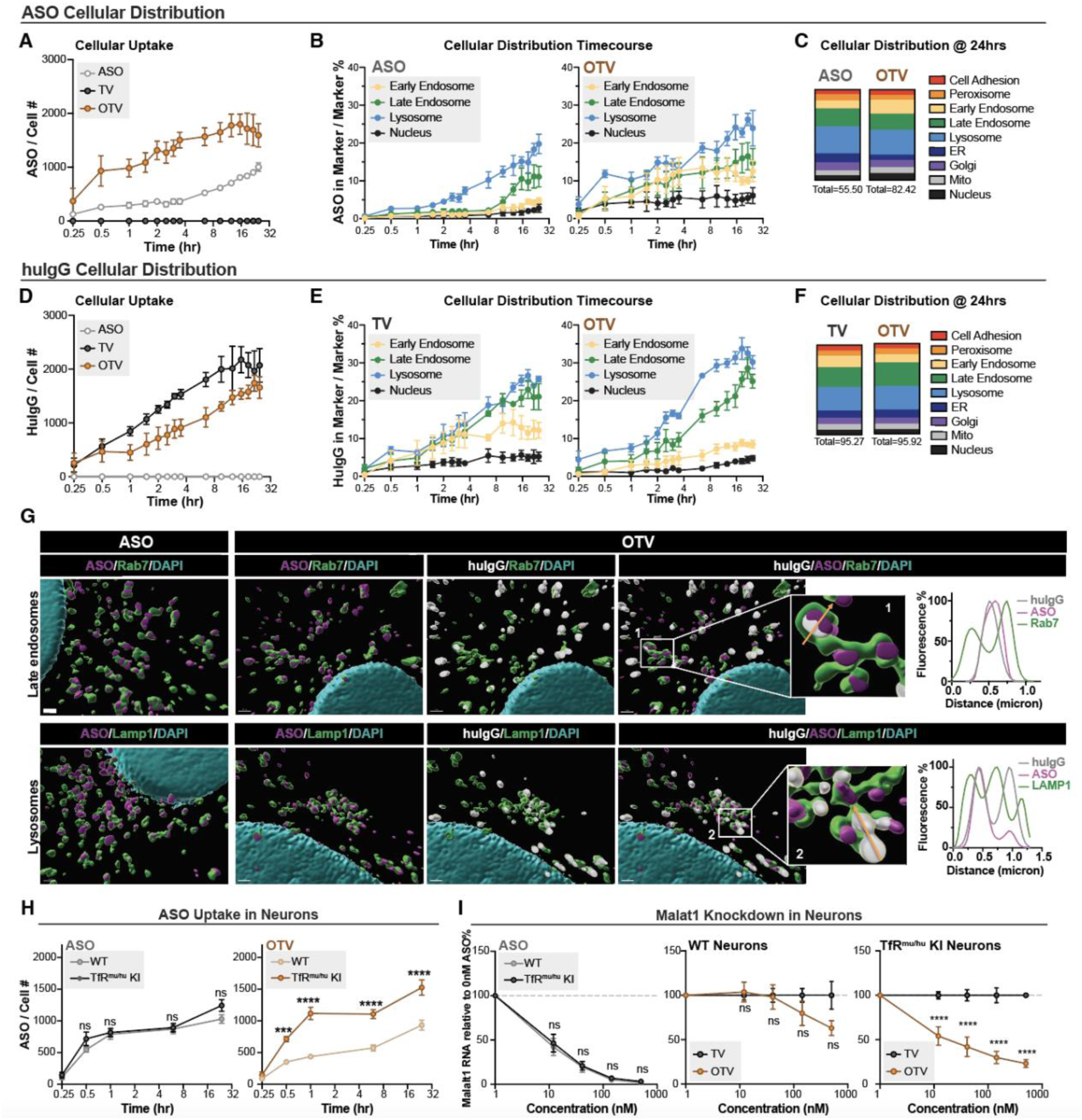
OTV drives cellular uptake via hTfR1 binding and does not alter ASO trafficking compared to naked ASO treatment. **(A-F)** Total ASO (A-C) and total human IgG (huIgG) (D-F) *in vitro* cellular trafficking following treatment with 40nM naked ASO, or the molar equivalent of OTV or TV. Live-cell total cellular uptake of ASO (A) and huIgG (D) into SH-SY5Y cells determined by measuring Cy-3 (ASO) or Alexa-647 (huIgG) fluorescence intensity. Live-cell accumulation over time of ASO (B) and huIgG (E) from naked ASO or OTV in indicated organelles along the main trafficking route using the same labeling as in (A) and (D). Total cellular distribution of ASO (C) and huIgG (F) at 24 hours. See Fig. S1 for time course of organelles not shown in (B) and (E). (**G**) Super resolution microscopy of anti-ASO (purple), anti-huIgG (white) and markers of late endosomes (Rab7) or lysosomes (Lamp1) (green) in SH-SY5Y cells treated with naked ASO or OTV for 24 hours prior to fixation and staining. Insets 1 and 2 show magnifications and associated line graphs of localization. (**H**) Uptake of 500nM ASO from naked ASO or OTV in wild-type (WT) or human TfR^mu/hu^ KI primary mouse neurons measured in fixed cells by anti-ASO staining. (**I**) *Malat1* levels measure by qPCR in WT or TfR^mu/hu^ KI primary mouse neurons following treatment with either naked ASO or OTV. Data are shown as means ± SEM; n = 3 independent experiments. Two-way ANOVA adjusted for multiple comparisons using Sidak’s method. ns p>0.05, ***p < 0.001, ****p < 0.0001

To understand the impact of ASO conjugation on target knockdown in an *in vitro* setting, we next compared uptake and RNA knockdown in primary neurons from TfR^mu/hu^ KI mice, a model that enables the characterization of human TfR-binding molecules in a physiological setting[23]. As the TV protein binds to human but not mouse TfR1, the presence of hTfR1 in TfR^mu/hu^ KI mouse primary neurons[23] led to substantially improved uptake of OTV compared to that observed in wildtype (WT) mouse primary neurons (Fig. 2H). Not surprisingly, primary neuron genotype had no effect on the uptake (Fig. 2H) or knockdown potency (Fig. 2I, left panel) of naked ASO. Consistent with the observation of poor OTV uptake in non hTfR1 expressing cells (Fig. 1D, right panel) and primary neurons (Fig. 2H), OTV was unable to knockdown *Malat1* RNA in WT neurons at lower concentrations (<100nM), though did achieve minimal knockdown at higher concentrations (>100nM) (Fig. 2I, middle panel). In TfR^mu/hu^ KI neurons, however, OTV treatment resulted in significant knockdown at all concentrations tested, demonstrating effective hTfR1-mediated uptake and target gene knockdown (Fig. 2I, right panel). Together, these *in vitro* studies show that while OTV is capable of providing hTfR1-mediated uptake of ASO, the subsequent subcellular trafficking of OTV-derived ASO is not significantly altered compared to naked ASO.

### Systemically Delivered OTV Knocks Down RNA Across CNS Regions and Cell Types In Vivo

Moving from an *in vitro* to an *in vivo* system, we next tested the ability of OTV to achieve *Malat1* knockdown in the CNS after peripheral dosing in mice. For the first *in vivo* assessment, we employed the constrained ethyl (cET) 2’sugar ASO chemistry following confirmation that both the cET chemistry and the locked nucleic acid (LNA) chemistry (used in all *in vitro* studies) provided robust and equivalent knockdown of endogenous *Malat1* RNA following CSF delivery in WT mice (Fig. S2A). Following a single IV dose of OTV, we first observed that conjugation to the TV protein enables a significant extension of ASO plasma half-life in WT mice (n=3/group; cohort 1) (Fig. 3A). Consistent with previous observations across species, naked ASO clears from circulation in minutes (Fig. 3A), largely depositing in the kidney and liver[35]. To next characterize OTV biodistribution *in vivo*, we dosed TfR^mu/hu^ KI mice (n=5/group) with either 1) a TV control without an ASO, 2) naked ASO, 3) ASO conjugated to a non-hTfR1 binding antibody (mAb:ASO), or 4) OTV at 2.4 mg/kg ASO or molar equivalence and collected tissues after one (cohort #2) and four (cohort #3) doses to compare ASO accumulation and pharmacodynamic response throughout the CNS and periphery (Fig. 3B). OTV deposited significantly more ASO in the brain and spinal cord compared to both naked ASO or mAb:ASO treatment groups (Fig. 3C, Fig. S2B). Further, OTV achieved significant ASO accumulation in the brain, depositing 9nM ASO after a single dose and 26nM after four weekly doses (Fig. 3C). Less than 10% of the ASO in brain remained conjugated to the TV 72 hours after the fourth dose (Fig. S2C), indicating robust ASO accumulation that persists well after TV deconjugation and/or degradation, though total huIgG is still readily detectable across CNS cells via huIgG staining in the hippocampus at 24hrs post-dose (Fig. S2D).

**Fig 3.**
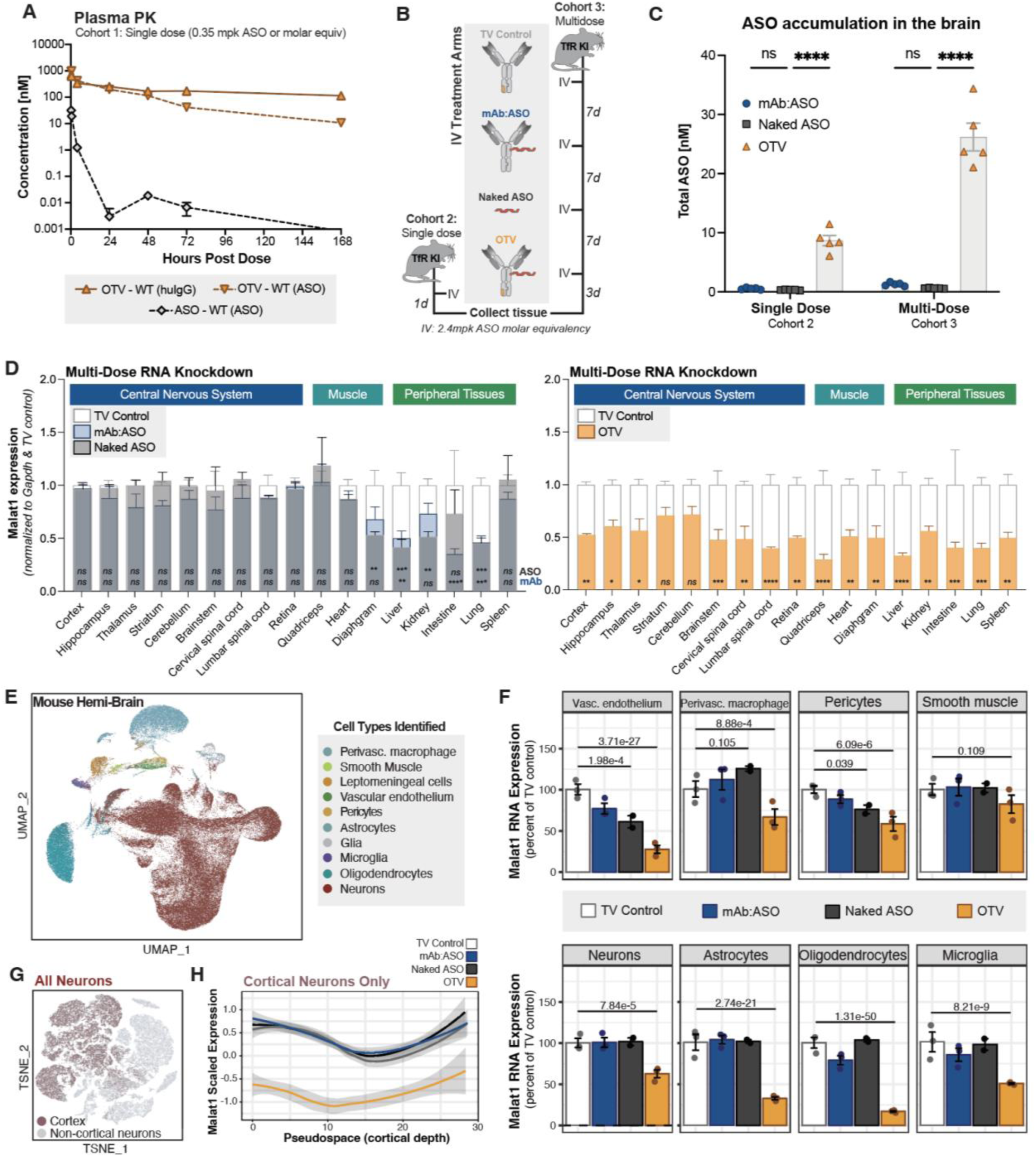
OTV enables ASO accumulation *in vivo* and achieves target RNA knockdown in CNS, muscle, and peripheral tissues, and in all major CNS cell types. **(A)** Pharmacokinetic time course of huIgG (solid line) and ASO (dashed line) in Cohort 1 plasma following intravenous delivery of 0.35mg/kg of naked ASO (grey), or the molar equivalent amount of OTV (orange) in WT mice, n=3-6 mice per group. (**B)** Study design schematic containing single and multi-dose arms corresponding to Cohort 2 and Cohort 3, respectively. (**C)** ASO concentration in the brain after one or four doses of OTV (orange), naked ASO (grey) or mAb:ASO (blue). (**D)** *Cohort 3: Malat1* RNA expression in various brain regions and peripheral tissues following four doses of a TV control with no ASO (white), OTV (orange), naked ASO (grey), and mAb:ASO (blue). Expression values are normalized to the housekeeping gene *Gapdh* and displayed relative to the TV control cohort set at a value of 1. In the left panel, top significance line corresponds to the ASO vs. TV comparison and bottom significance line corresponds to the mAb:ASO vs. TV comparison. (**E)** *Cohort 3:* UMAP dimensionality reduction showing all nuclei passing QC thresholds. Cell type predictions were assigned using the Mouse Brain Atlas adolescent mouse scRNA-seq data set as a reference (Zeisel et al, 2018). Nuclei labeled “glia” could not be assigned a more granular cell type. (**F)** Boxplots showing pseudobulked scaled expression values for Malat1 across cell types. Each dot represents one animal. **(G)** TSNE dimensionality reduction showing a subclustering of all telencephalon neurons. **(H)** Local regression plots showing moving average *Malat1.* Data are shown as means ± SEM. One-way ANOVA adjusted for multiple comparisons using Dunnett’s method for (C); Two-way ANOVA adjusted for multiple comparisons to the TV control group using Dunnett’s method for (D). ns p>0.05, * p<0.05, ** p<0.01, *** p< 0.001 **** p< 0.0001.

The ubiquitous expression of *Malat1* allows for pharmacodynamic interrogation throughout the body. We examined ASO accumulation and *Malat1* RNA expression in various CNS and peripheral tissues, focusing on therapeutically relevant regions and those that are difficult to reach with naked ASOs, including deep brain structures (e.g., striatum)[36] and muscle tissues (e.g., quadricep)[3, 37]. OTV resulted in significantly more robust *Malat1* RNA knockdown throughout the brain and spinal cord compared to naked ASO or mAb:ASO, consistent with differences in ASO deposition across these groups (Fig. 3D). OTV also resulted in significantly greater *Malat1* knockdown in the quadricep and heart tissues. Interestingly, the degree of knockdown in the quadricep and heart was significantly greater after OTV administration compared to either naked ASO or mAb:ASO, despite similar levels of ASO accumulation (Fig. S2B), suggesting that hTfR1-mediated uptake might direct the ASO toward a more functional intracellular compartment. We observed similar levels of *Malat1* knockdown across groups in tissues involved in peripheral clearance of ASOs (liver and kidney), indicating similar knockdown dynamics in these tissues despite the extended ASO half-life afforded by conjugation to the TV protein.

To examine the capacity of OTV to knockdown RNA across cell types in the brain, we utilized single-nucleus RNA-sequencing (snRNA-seq). Following homogenization of brain tissues, nuclei were isolated from mouse hemibrains across the four treatment groups: TV Control (n = 3), mAb:ASO (n = 3), naked ASO (n = 2), and OTV (n = 3). Predictive label transfer using the Mouse Brain Atlas as a reference[38] uncovered expected major cell types which were validated by expression of canonical marker genes (Fig. 3E; Fig. S2E, F). To assess *Malat1* knockdown, pseudobulk counts were generated for each major cell type and differential expression was carried out using DESeq2[39]. Strikingly, mice treated with OTV showed significant knockdown of *Malat1* RNA compared to vehicle treatment in nearly all cell types, including astrocytes, oligodendrocytes, neurons, microglia, vascular smooth muscle, vascular endothelium, pericytes, and perivascular macrophages (Fig. 3F). In contrast, naked ASO only achieved knockdown in pericytes and vascular endothelium.

Next, we examined whether target knockdown in neurons remained consistent across all layers of the cortex. Cortical neurons were identified by iterative subclustering and predictive labeling (Fig. 3G; Fig. S2G-I). To predict cortical depth as a function of gene expression, UMAP dimensionality reduction was carried out using a subset of genes that are known to be patterned across layers of the cortex[40, 41]. Pseudotime was then calculated to approximate cortical depth. The resulting pseudospace trajectory accurately predicted expected patterns in cortical layer markers (Fig. S2J). *Malat1* expression was substantially lower in OTV-treated cortical neurons across the entire pseudospace trajectory, with no appreciable difference in knockdown at deeper levels (Fig. 3H). These data demonstrate hTfR1 mediated uptake of OTV into the CNS following IV administration in TfR^mu/hu^ KI mice that subsequently resulted in target RNA knockdown across all CNS cell types, including uniform knockdown in neurons across cortical layers.

### OTV Differentiates from High Affinity, Bivalent TfR:ASO Drug Conjugate in the CNS

OTV was engineered to bind hTfR1 in a monovalent format—distinct from existing bivalent transferrin receptor antibodies that engage two transferrin receptors simultaneously [42]. To understand the impact of bivalent, high affinity hTfR1 binding and how that may differentiate from OTV, we developed a protein-ASO conjugate, termed TfR:ASO (Fig. 4A), using a bivalent antibody with a higher affinity to hTfR1 (0.12 nM). As with OTV, the addition of an ASO does not impact hTfR1 binding (Fig. S3A-C). The binding affinity of the TfR:ASO molecule to cynomolgus monkey TfR1 is much weaker (86 nM; Fig. S3B,C), indicating that data collected in this species may not be as translatable to humans. For this reason, we performed all *in vivo* studies comparing TfR:ASO to OTV in the TfR^mu/hu^ KI mouse model and used LNA rather than cET ASO chemistry due to ease of molecule generation.

**Fig 4.**
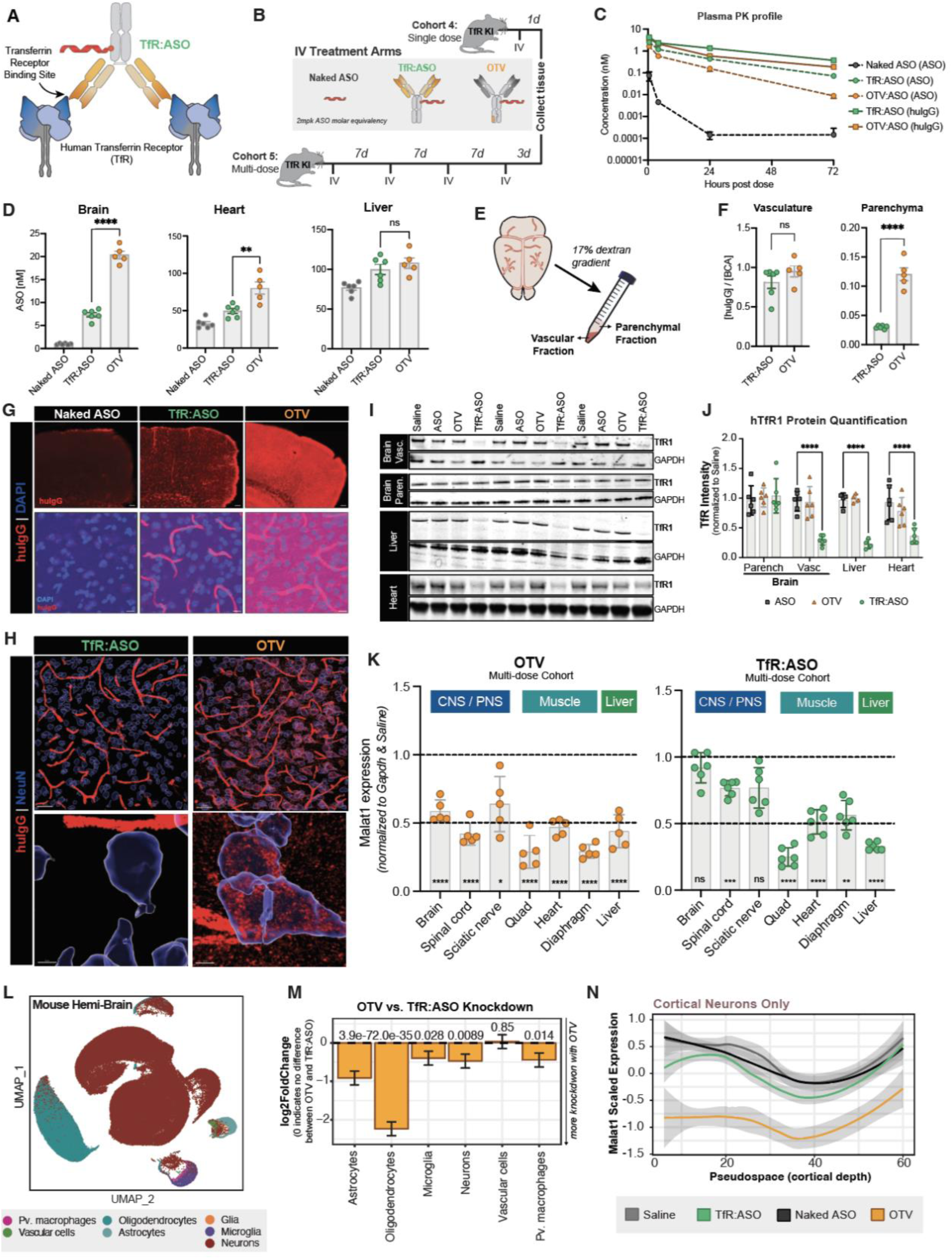
A high affinity, bivalent TfR:ASO conjugate degrades TfR and results in limited ASO deposition in the CNS. **(A)** Schematic of bivalent TfR binding. (**B)** Study design schematic containing single and multi-dose arms corresponding to Cohort 4 and Cohort 5 respectively. **(C)** Pharmacokinetic profile of ASO concentration (solid lines) and huIgG concentration (dashed lines) after a single dose of naked ASO (grey), TfR:ASO (green), or OTV (orange). (**D)** *Cohort 5*: ASO concentration in the brain, liver, and heart after treatment with either naked ASO (grey), TfR:ASO (green), or OTV (orange). **(E)** Schematic of capillary depletion assay to separate brain vasculature from parenchyma. (**F)** *Cohort 4:* huIgG quantification in vasculature and parenchymal fractions after capillary depletion. **(G)** *Cohort 4:* Immunofluorescent staining of huIgG in cortical tissue section from TfR^mu/hu^ KI mice dosed with either naked ASO, OTV, or TfR:ASO. Scale bars in top panels = 100µm; scale bars in bottom panels = 10µm. (**H)** Higher resolution view of (G) after generating a surface around all NeuN+ neurons using Imaris software. Scale bars in top panels = 30µm; scale bars in bottom panels = 4µm. (**I)** *Cohort 5:* Western blot displaying levels of transferrin receptor in the brain vasculature, brain parenchyma, liver, and heart after dosing with Saline, naked ASO, OTV, or TfR:ASO. (**J)** Quantification of western blot images in (I). (**K)** *Cohort 5*: *Malat1* knockdown in various tissues after OTV treatment (left) and TfR:ASO treatment (right), normalized to *Gapdh* expression and displayed as a fold change relative to Saline-treated mice. (**L)** *Cohort 5:* UMAP dimensionality reduction showing all nuclei passing QC thresholds with assigned cell type predictions. (**M)** Barplots showing differential expression for *Malat1* in pseudobulked data between OTV and TfR:ASO treatments. Bars indicate Log^2^ FoldChange in OTV compared to TfR:ASO across all cell types. (**N)** Local regression plots showing moving average *Malat1* scaled expression in each treatment group across predicted cortical depth calculated with 500 randomly selected cells from each condition. Shaded regions show 95% confidence interval. Data are shown as Mean ± SEM. Two-way ANOVA adjusted for multiple comparisons to the Saline group using Dunnett’s method. ns p>0.05, * p<0.05, ** p<0.01, ***p< 0.001, ****p< 0.0001.

We first compared plasma pharmacokinetics of OTV and TfR:ASO in TfR^mu/hu^ KI mice (Fig. 4B) and observed faster clearance of OTV in plasma relative to the TfR:ASO high affinity, bivalent molecule (Fig. 4C) (n=5-6/group). Interestingly, when incubated with full length hTfR1 protein OTV displays a single narrow peak via dynamic light scattering, while TfR:ASO displays a very broad peak, indicating aggregation of the complex *ex vivo* specifically when engaging hTfR1 (Fig. S3D). Dimerization and aggregation of the TfR:ASO-hTfR1 complex *in vivo* may therefore explain the slower clearance of TfR:ASO despite increased binding affinity to hTfR1, as it is known that when hTfR1 dimerizes, it forces internalization and degradation of the receptor itself[43].

Next, to identify differences in biodistribution between OTV and TfR:ASO, we measured ASO concentration in the brain, heart, and liver after four 2mg/kg ASO equivalent doses of each molecule (Cohort 4; n=5-6/group). OTV deposited significantly more ASO in the brain and heart, but comparable amounts in liver as TfR:ASO, despite the ∼1000X difference in hTfR1 binding affinity (Fig. 4D). A potential explanation for the significant difference in ASO accumulation in CNS and muscle tissues is the disparity in hTfR1 affinity and valency between the two molecules. Crossing the TfR1-expressing endothelial cell layer at the BBB [44] is a requirement for TV deposition in the brain parenchyma. We hypothesized that this high affinity, bivalent TfR:ASO molecule might result in reduced transcytosis and vascular trapping [21, 27, 45–47], thus accumulating drug specifically in brain vasculature and not parenchymal tissue. To test this hypothesis we stained for huIgG 24 hours after a single dose of OTV or TfR:ASO and observed widespread, homogenous huIgG signal throughout the brains of OTV-dosed mice, but very heterogenous distribution in TfR:ASO-dosed mice, with robust huIgG signal in vasculature but minimal huIgG detectable in parenchymal tissue (n=4/group) (Fig. 4G). There was almost no overlap with NeuN in mice treated with TfR:ASO (Fig. 4H), indicating very little transcytosis into the parenchyma of the brain. This striking difference was confirmed by huIgG ELISA after brain capillary depletion, which showed significantly more huIgG in the parenchyma of mice dosed with OTV, but comparable huIgG levels in the vasculature of OTV- and TfR:ASO-dosed mice (Fig. 4E-F).

A possible mechanism underlying the pharmacokinetic and distribution differences between OTV and TfR:ASO could be the internalization and degradation of hTfR1 upon bivalent binding [43, 48]. To investigate this, we measured hTfR1 protein levels in various tissues to determine if TfR:ASO resulted in loss of hTfR1 *in vivo*. Strikingly, hTfR1 protein levels were significantly reduced in the liver, heart, and brain vasculature, but not brain parenchyma (Fig. 4I-J; Fig. S3E), only in the TfR:ASO treated group, providing a valid explanation for a) slower systemic clearance despite a higher affinity to TfR1, and b) limited BBB crossing along with significantly restricted distribution of TfR:ASO compared to OTV.

To further assess the impact of differences in biodistribution on target knockdown, we measured *Malat1* RNA expression in various tissues after four doses of either OTV, TfR:ASO, or naked ASO. As anticipated, peripherally delivered ASO only achieves robust knockdown in the liver (Fig. S3F), with limited target engagement in tissues protected by the BBB. OTV achieves superior knockdown in both the brain and spinal cord compared to TfR:ASO (Fig. 4K), consistent with greater ASO deposition in the CNS parenchyma. OTV also provides significantly more knockdown in the diaphragm compared to TfR:ASO. Interestingly, we observed equivalent knockdown in the liver, quadricep muscle, and heart, despite differences in hTfR1 affinity.

To compare target knockdown resulting from OTV versus TfR:ASO across all cell types in the brain, we again employed snRNA-seq. Single nuclei suspensions were isolated from mouse hemibrains across the four treatment groups: vehicle control (n = 4), naked ASO delivered systemically (n = 4), TfR:ASO (n = 6), and OTV (n = 3). As described previously, major cell types were identified through predictive labeling and marker gene analysis (Fig. 4L; Fig. S3G). OTV again achieved knockdown across nearly all major cell types (Fig. S3H). While TfR:ASO did induce some knockdown in astrocytes and oligodendrocytes, OTV induced significantly greater knockdown in these cell types (Fig. 4M; Fig. S3H). Notably, OTV uniquely induced broad knockdown in neurons compared with either naked ASO or TfR:ASO, consistent with imaging results (Fig. 4H). Pseudospace analysis in cortical neurons revealed that this difference is most pronounced at deeper cortical levels (Fig. S3I). TfR:ASO induces a slight decrease in *Malat1* levels in the more superficial layers of the cortex, but this difference is entirely abrogated in the deepest layers. In contrast, OTV again induced knockdown uniformly across all cortical layers (Fig. 4N).

### OTV Preserves Long ASO Half-life and Knockdown Efficiency in vivo

Current ASO therapies targeting CNS disorders primarily rely on intrathecal (IT) delivery to administer ASOs directly to the CSF. Direct CNS delivery of ASOs in rodents, however, is primarily accomplished via intracerebroventricular (ICV) injection into the lateral ventricle. ASO distribution in the brain following either of these methods is significantly impacted by a number of factors, including infusion volume and speed [49]. By comparison, OTV delivers ASO to the brain via transcytosis through the extensive capillary network comprising the BBB. Despite these two different routes, we hypothesized that the rate of ASO metabolism in mouse tissue would be similar between both methods of delivery as naked ASO and OTV showed similar subcellular distribution patterns *in vitro* (Fig. 2). To compare ASO half-life and knockdown duration of action, we measured ASO concentration and *Malat1* RNA expression in the brain, spinal cord, and liver at several timepoints after IV OTV loading administration (six doses of 0.9mg/kg ASO molar equivalence; Cohort 6) or ICV ASO administration (50µg, equivalent to ∼one dose of 2.4mg/kg ASO; Cohort 7). In both cohorts of mice, ASO half-life and knockdown duration of action were longest in the brain and shortest in the liver, which is favorable for a neurotherapeutic agent (Fig. 5A-B; Fig. S4A-D). Importantly, ASO half-life was comparable between routes of delivery in all tissues (Fig. 5B; Fig. S4C), suggesting that the TV does not negatively impact ASO stability once delivered to cells *in vivo*.

**Fig 5.**
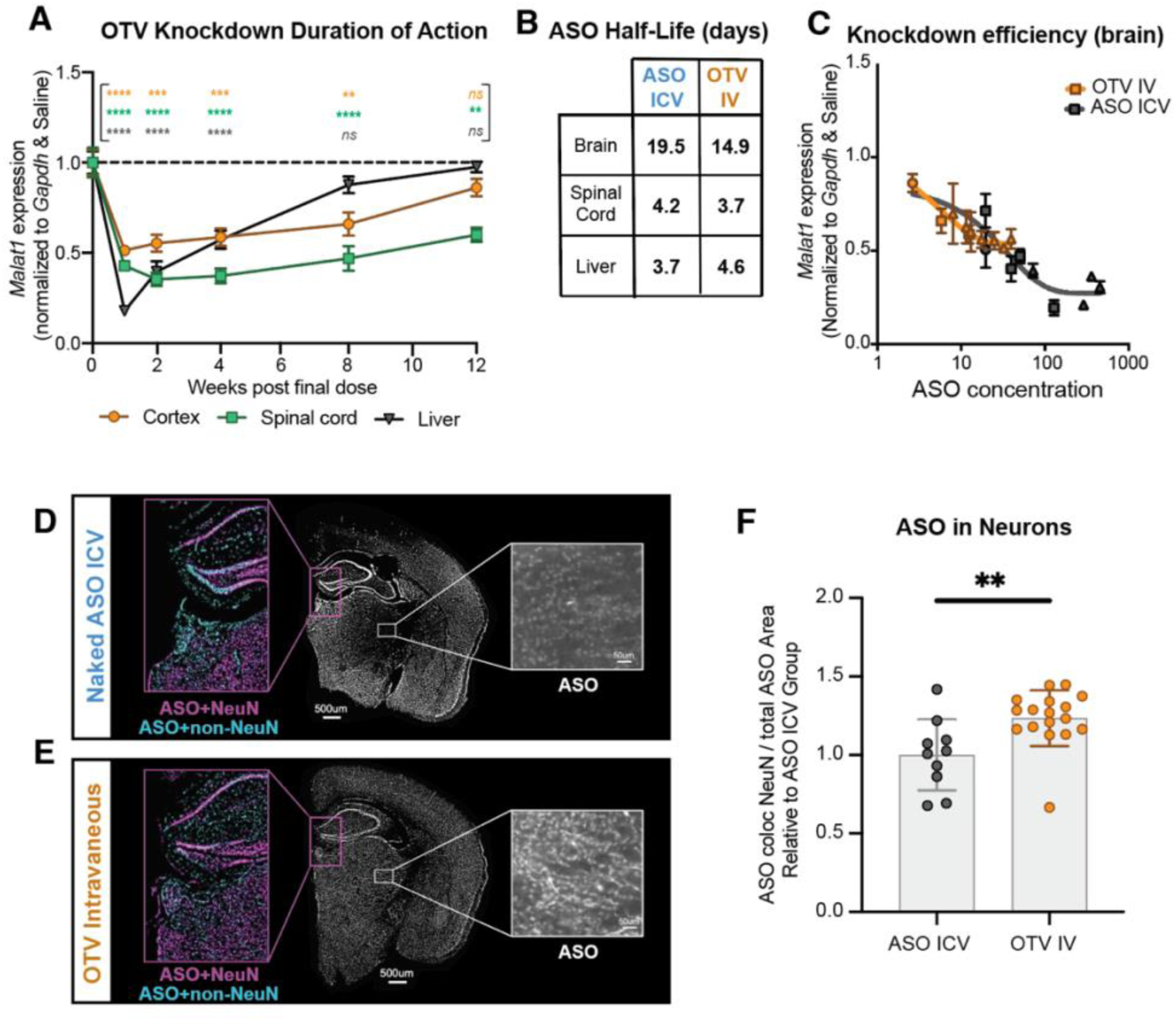
OTV preserves long half-life and knockdown efficiency in the mouse brain. **(A)** *Malat1* expression in the mouse brain (orange), spinal cord (green), and liver (grey) at various timepoints after the last loading dose of OTV. (**B)** ASO half-life in days in the mouse brain, spinal cord, liver, and kidney after OTV or ICV administration. Data in Fig. S4A,B were used to calculate the ASO half-life. (**C**) Brain ASO concentration plotted against *Malat1* RNA expression as a measure of knockdown efficiency. Triangles represent collection timepoints under 2 weeks, squares represent 4- and 8-week collection timepoints, circles represent 12- and 14-week collection timepoints. (**D-E)** Immunofluorescent staining of mouse brain tissue with an anti-ASO antibody after ICV (D) or OTV (E) administration at 1 week post-dose. Middle panel = a coronal hemibrain section; Right panel = zoomed in view of the thalamus; Left panel = colocalization with NeuN (purple) or non-neuronal DAPI (turquoise). **(F)** Quantification of colocalization between ASO and NeuN signal, normalized to the ASO ICV cohort. Data are shown as means ± SEM. Two-way ANOVA adjusted for multiple comparisons to the Saline-dosed group using Dunnett’s method in (A). Student’s t-test performed between the ASO ICV and OTV group at 1 week in (F). ns p>0.05, * p<0.05, ** p<0.01, ***p< 0.001 ****p< 0.0001

We next sought to understand the efficiency of target knockdown from OTV versus naked ASO. The dose-response curves for OTV and naked ASO in the mouse brain are overlapping (Fig. 5C), suggesting equivalent knockdown potency for the ubiquitously expressed target gene, *Malat1*. Thus, conjugation to a TV does not inhibit knockdown potency in the brain. Despite direct CSF delivery during ICV injection, a significant percentage of the ASO delivered enters systemic circulation and deposits in peripheral tissues[50]. As both OTV and naked ASO are known to accumulate in the liver, we also generated dose response curves to compare knockdown efficiency in the liver. Interestingly, the knockdown efficiency of naked ASO is greater than OTV in the liver (Fig. S4E), suggesting potential differences in ASO distribution, trafficking, and/or metabolism in this tissue depending on the mechanism of ASO uptake.

Finally, we stained mouse brain sections with an anti-ASO antibody developed against the chemically modified phosphorothioate backbone to compare ASO biodistribution after OTV or ICV delivery. While a bolus of ASO delivered directly to the lateral ventricle can deposit a large amount of ASO to the mouse brain, we observed that subcortical regions furthest from CSF circulation, such as the thalamus, display less ASO accumulation (Fig. 5D-E). Roughly 30-40% of the ASO signal colocalizes with neuronal cell bodies (NeuN+), indicating substantial ASO uptake into neurons in both conditions. Interestingly, similar to the uptake we observed in SH-SY5Y cells (Fig. 2), ASO delivered by OTV showed a modest but significant increase in neuronal colocalization compared to CSF delivered naked ASO (Fig. 5F), suggesting that hTfR1 binding may promote neuronal uptake, as anticipated based on TfR1 expression patterns in the brain [51, 52].

### OTV Drives Significant and Cumulative ASO Uptake in the Non-Human Primate CNS

ASO biodistribution via direct CSF delivery is expected to vary between rodents and primates. First, biophysical considerations and experimental data on diffusional transport of macromolecules from CSF into adjacent brain tissue predict similar penetration distances across species, thus larger brains with longer diffusion distances may have more obvious surface gradients and less efficient delivery to deeper brain structures [53]. Further, the site of administration in mice (lateral ventricle) is more proximal to many brain structures compared to the lumbar spinal cord, the most common injection site for both NHPs and humans. Finally, the ratio of typical dose volume to CSF volume is much greater for rodent ICV injections compared to intrathecal injections [54]. In order to further understand ASO biodistribution in a primate species, we selected the Cynomolgus – *Macaca fascicularis* – species of NHPs for our studies. To that end, we generated an OTV with improved binding affinity to Cynomolgus TfR1 (Fig. S5A) and conjugated an ASO targeting Cynomolgus monkey *MALAT1,* termed ASO^Mf^. After confirming cellular uptake and knockdown functionality in Cynomolgus fibroblasts *in vitro* (Fig. S5B-C), we dosed n=3 Cynomolgus monkeys with a 4mg IT dose of naked ASO^Mf^ to mimic the current standard of care for ASO therapies with a 2 week post dose tissue collection timepoint (Fig. 6A). A 4mg dose was selected as it was the highest tolerated dose of naked ASO^Mf^ administered intrathecally. Additionally, allometric scaling results in an average dose of 1.2 mg/kg, which falls within the typical range of clinically dosed ASOs in human. In the same study for proper comparison, we administered a short burst of loading IV doses of OTV or naked ASO^Mf^ at 1.1mg/kg ASO molar equivalence and collected tissues 2 weeks post the last dose (Fig. 6A). Importantly, we selected IT ASO^Mf^ and IV OTV doses to exposure match CNS ASO concentrations based on anticipated ASO retention rates in the CNS [35, 53], and dosed the molar equivalent amount of ASO^Mf^ intravenously to serve as a TfR1 uptake control group. Of note, all three monkeys in only the intrathecal group experienced adverse events at dosing (e.g., hindlimb paralysis).

**Fig 6.**
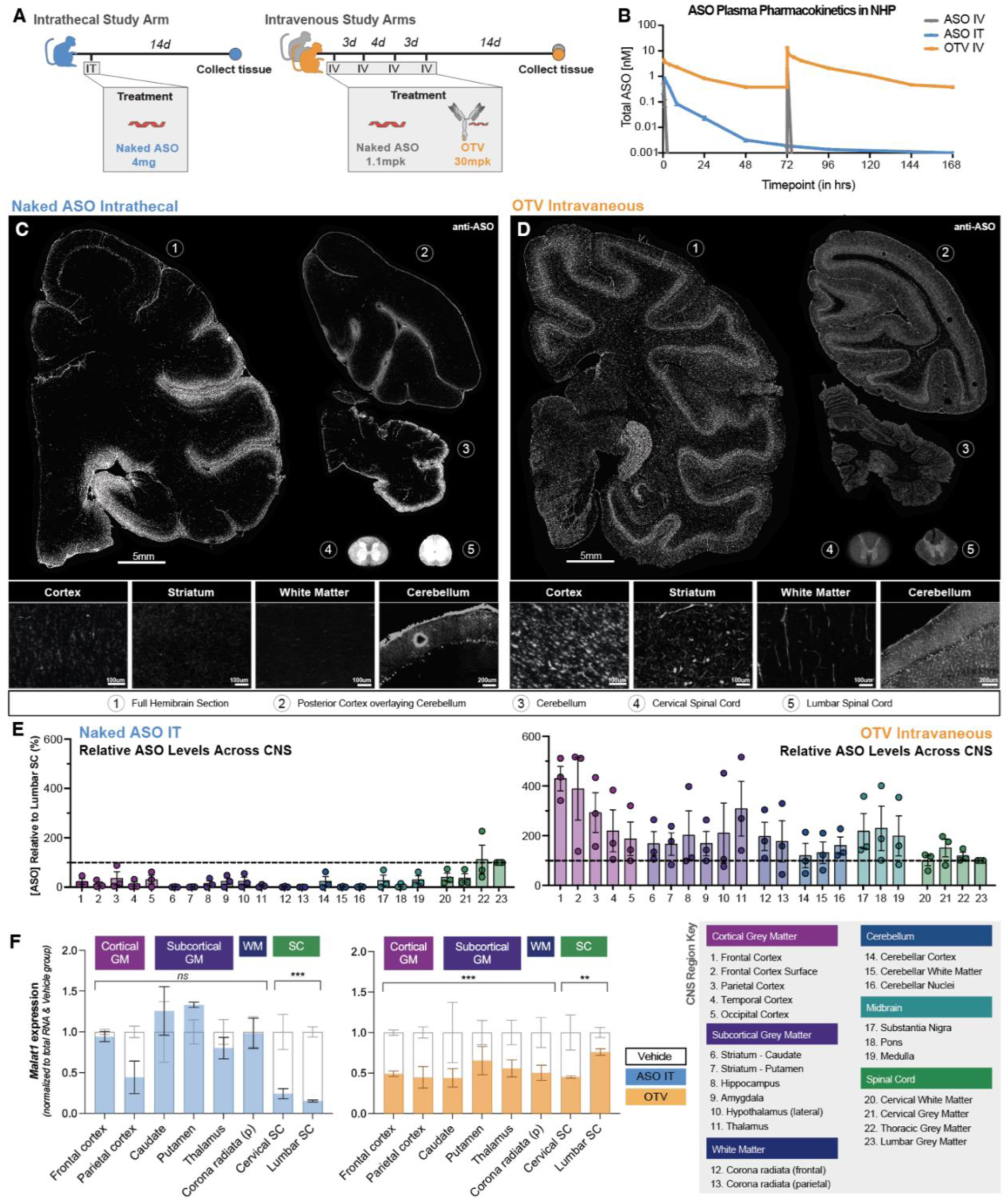
OTV drives widespread ASO^Mf^ accumulation and target knockdown in the Non-Human Primate CNS compared to IT dosing of naked ASO. (A) NHP study design schematic. (B) Plasma ASO pharmacokinetic profile following intravenous dosing of naked ASO^Mf^ (grey), intravenous dosing of OTV (orange), and intrathecal dosing of naked ASO^Mf^ (blue). (C) Top: Hemibrain and spinal cord sections from a cynomolgus monkey treated with naked ASO dosed intrathecally; stained with an anti-ASO antibody (white). Bottom: Zoomed in images of cortex, striatum, cerebellum, and white matter. (D) Top: Hemibrain from a cynomolgus monkey treated with OTV dosed intravenously; stained with an anti-ASO antibody (white), Bottom: Zoomed in images of cortex, striatum, cerebellum, and white matter. (E) Quantification of total ASO^Mf^ concentration in 23 regions throughout the brain and spinal cord (ASO^Mf^ IT – left; OTV IV – right). Data presented relative to concentration in lumbar spinal cord, the site of IT injection. Key of CNS regions below. (F) *MALAT1* RNA knockdown in eight regions throughout the brain and spinal cord, normalized to total RNA input and displayed as fold change relative to vehicle-treated cynomolgus monkeys. Data are shown as means ± SEM. Two-way ANOVA for brain and spinal cord regions treated separately, with significance values displayed for the effect of Cohort on knockdown in brain and spinal cord. ns p>0.05, ***p< 0.001 ****p< 0.0001

As anticipated, OTV provided significant ASO half-life extension compared to naked ASO (Fig. 6B; Table S1). In this higher species, we expanded our peripheral tissue assessment to include other tissues that have high physiological iron demand. In addition to the tissues assessed in mice (brain, spinal cord, quadricep muscle, liver and kidney), we investigated ASO^Mf^ levels in the diaphragm, retina, and bone marrow (Fig. S6A-I). OTV provided enhanced exposure compared to naked ASO^Mf^ in all tissues except the kidney. This benefit was especially noticeable in the retina (Fig. S6G), where ASO^Mf^ concentration is undetectable after delivery of naked ASO^Mf^. Notably, ASO^Mf^ delivery by OTV was significantly higher across muscle tissues, consistent with enhanced ASO staining and *MALAT1* RNA knockdown in muscle tissue after OTV administration (Fig. S6M, P, U-W).

To characterize ASO^Mf^ distribution throughout the CNS, we stained brain and spinal cord sections with an ASO-specific antibody. As anticipated, naked ASO^Mf^ delivered intravenously is absent from the CNS, as it clears quickly and does not effectively cross the BBB (Fig. S7A). IT delivery results in heterogenous ASO^Mf^ distribution throughout the CNS, with a striking gradient from lumbar spinal cord to the brain (Fig. 6C). By contrast, OTV deposits ASO^Mf^ evenly throughout the brain and spinal cord, leading to homogenous biodistribution (Fig. 6D). After IT delivery, ASO^Mf^ accumulation is much greater ventrally, as well as near the surface of the brain. Deep brain structures, including the thalamus, appear nearly devoid of ASO^Mf^. After OTV administration, however, we observe greater ASO^Mf^ accumulation in regions that are traditionally challenging to reach with IT injection, such as the striatum, cerebellum, and white matter (Fig. 6C-D). Further, IT delivery results in significantly more variability in ASO^Mf^ distribution throughout the brain, especially in cortical areas where some regions display high ASO accumulation and others show very little (Fig. S7B-C). To confirm these imaging results, we dissected 23 regions of the brain and spinal cord (Table S2) and measured ASO^Mf^ concentration in each. As anticipated, ASO^Mf^ accumulates in the brain with additional doses of OTV (Fig. S7D), allowing our OTV dosing regimen to achieve similar levels of total ASO^Mf^ in the CNS as IT dosing (Table S3). Consistent with the imaging results, systemic administration of OTV results in significantly more homogenous biodistribution of ASO^Mf^ throughout the CNS (Fig. 6E). After IT delivery, ASO^Mf^ concentration is substantially higher in the lumbar spinal cord (i.e., near the site of injection) compared to the brain (6-fold higher on average). Even cervical spinal cord, which is closer to the injection site than the brain, exhibits over 60% less ASO^Mf^ accumulation than the lumbar spinal cord, highlighting the declining caudal-to-rostral distribution gradient that often arises with IT delivery of macromolecules to the lumbar CSF space [55]. In the brain, ASO^Mf^ concentration is unevenly distributed after IT delivery, with relatively low amounts deposited in deep structures such as the striatum. By contrast, after systemic OTV administration, ASO^Mf^ concentration is comparable between the spinal cord and all brain regions sampled, including deep brain structures.

These differences in ASO^Mf^ biodistribution culminate in substantially less target knockdown in the brain versus spinal cord following IT administration, compared to more uniform levels of knockdown across CNS regions following systemic OTV administration (Fig. 6F). Specifically, after IT dosing we observe substantial target knockdown in the spinal cord, no target knockdown in subcortical grey matter, and variable knockdown in cortical regions throughout the brain, consistent with ASO^Mf^ staining and exposures (Fig. S7E). Unlike IT dosing, OTV results in ∼50% knockdown across various brain regions, making it more desirable for indications that affect the brain globally.

Given the much larger scale of the Cynomolgus monkey brain along with the observed differences in ASO^Mf^ distribution after OTV vs. IT dosing, we sought to further explore the capability of OTV to reach and deliver efficient knockdown to deep brain regions. To this end, we employed snRNA-seq on dissected thalamus tissue from cynomolgus monkeys treated with naked ASO^Mf^ or OTV (n = 3 for each). We recovered major CNS cell types: a large population of oligodendrocytes and smaller populations of oligodendrocyte precursor cells (OPCs), astrocytes, neurons, and microglia (Fig. S8A-B). Interestingly, compared to vehicle-treated monkeys we observed modest knockdown of *MALAT1* after IT injection in oligodendrocytes and microglia, with *MALAT1* expression only trending lower in other cell types. However, OTV induced stronger and more consistent knockdown in all cell types (Fig. S8C), confirming the superior ability of OTV to target deep structures in the NHP brain.

## Discussion

Advances in understanding the genetic and molecular basis of neurodegenerative disorders have enabled clinical interrogation of gene expression changes to modify disease progression. Toward this end, ASOs have been successfully used to treat disorders such as spinal muscular atrophy [56], but have failed to provide clinical benefit in others, such as Huntington’s disease [57]. Indeed, ASOs cannot cross the BBB on their own and must therefore be delivered to the CNS directly, either by lumbar puncture to the CSF surrounding the spinal cord or via ICV infusion [58]. These methods of delivery can be associated with increased adverse events (e.g. hindlimb paralysis observed at 4mg IT dose of ASO^Mf^ in NHPs) and result in limited biodistribution throughout the CNS. Here we engineered a novel platform, OTV, to transport ASOs across the BBB after systemic administration. By conjugating an ASO to a hTfR1-binding TV, we enable significant and cumulative ASO deposition throughout the brain and spinal cord, as well as in many peripheral tissues, including skeletal muscles and nerves. Enhanced ASO deposition to the CNS is accompanied by target knockdown across brain regions and brain cell types. Importantly, we use snRNAseq to demonstrate target knockdown in all major cell types in cynomolgus monkey brain tissue. This represents a meaningful advance in systemic ASO delivery by broadening the potential indication space to include CNS and neuromuscular disorders, as well as peripheral neuropathies.

Herein, we directly compare OTV to two alternative delivery strategies being pursued in the field: 1) ASO conjugated to a high affinity, bivalent anti-hTR1 antibody, and 2) naked ASO administered intrathecally. We demonstrate that OTV results in significantly more widespread ASO deposition throughout the CNS compared to both alternative therapeutic strategies. While high affinity, bivalent antibodies also engage hTfR1 along the brain’s vasculature, they cannot effectively cross the BBB into the parenchyma, highlighting the unique transcytosis properties of the TV. Recent studies in wildtype mice have proposed the use of antibodies directed against mouse TfR1 to deliver ASOs to the brain [42]; however, they show minimal to no improvement in ASO target engagement in the CNS and muscle above non-TfR1-binding antibody controls. This published approach likely suffers the same limitations as the TfR:ASO molecule used in the current study due to similar binding properties that restrict CNS biodistribution. Our results provide two potential explanations for the lack of added efficacy above a non-TfR1-binding antibody control. First, the high affinity binding of this bivalent molecule to TfR1 results in significant degradation of TfR1 protein, which prevents further receptor engagement, limits biodistribution, and poses a potential safety liability. Additionally, high affinity TfR1 binding of TfR:ASO leads to inferior CNS biodistribution as the molecule remains associated with endothelial cells rather than being released in the brain parenchyma, similar to what has been shown in previous work with lysosomal enzymes conjugated to TfR1 binding antibodies [27].

Intrathecal ASO injections can enable delivery directly to the CNS, relying in part on convection through CSF to reach the cranial region with subsequent transport into the brain via diffusion and/or perivascular transport [49]. However, diffusion of macromolecules from CSF into brain is limited and does not scale with brain size[53]. This results in an ASO gradient that does not effectively reach deep brain structures, as was observed in our NHP study. This sharp concentration gradient following intrathecal delivery results in regions with both extremely high (spinal cord) and low (striatum) ASO deposition, leading to variable knockdown throughout the CNS. This variability makes achieving a targeted level of knockdown required for efficacy (e.g. 40-60% [59]) in hard to reach brain regions much more difficult as there is often a concurrent desire to prevent toxicity that can result from very high local ASO concentrations and/or complete loss of function of the target gene in more CSF accessible CNS regions. Because OTV is delivered evenly via the extensive vasculature of the CNS, target knockdown is more precisely titratable. Additionally, we observe that roughly the same proportion of injected ASO remains in the CNS after OTV or IT dosing regimens using comparable concentrations of ASO (Table S3), and OTV dosing results in a similarly prolonged knockdown of gene expression in brain.

While the studies reported herein provide compelling proof of concept evidence for an OTV platform *in vivo*, there are numerous follow-up studies that will be essential to better understand the implications of altering dose level and dose frequency with an eye towards clinical translation. Additionally, more work is needed to understand the level of modularity of OTV as it pertains to protein architecture, other ASO targets, and other oligonucleotide formats, such as siRNAs. One area not addressed in this manuscript is the safety assessment of deposited ASO in peripheral clearance organs, such as liver and kidney. As ASO toxicity profiles have been shown to be sequence dependent [60], these endpoints are best examined in future OTV studies with therapeutic oligonucleotides. Finally, as is also the case for direct CSF administration of ASO [35] and existing ASO-conjugate technologies [15], OTV administration results in ASO being deposited in peripheral clearance organs in addition to the CNS; thus, strategies to further improve CNS targeting are warranted.

In sum, OTV has the potential to be highly versatile platform for efficient delivery and biodistribution of oligonucleotides to the CNS and other refractory tissues. ASOs can be generated against any gene product, thus enabling single- or multi-target approaches to address targets that are difficult to drug with other therapeutic modalities, such as antibodies or small molecules. Our novel OTV approach shows promise in both rodents and non-human primates, highlighting its potential for the effective treatment of neurological and neurodegenerative disorders.

## Materials and Methods

### Antisense Oligonucleotide Sequences

Sequences described for mouse and NHP where “#” denotes a constrained ethyl (cET) and “+” denotes a locked nucleic acid (LNA) to a subsequent nucleotide base and “*” denotes a phosphorothioate backbone linkage. All cytosines are methylated. The exact mouse *Malat1* ASO sequences [29] and NHP *MALAT1* sequence [30] are found below:

*Mouse Malat1 ASO Sequence with cET containing chemistry: 5’*

*#G*#C*#A*T*T*C*T*A*A*T*A*G*C*#A*#G*#C 3’*

*Mouse Malat1 ASO Sequence with LNA containing chemistry: 5’*

*+G*+C*+A*T*T*C*T*A*A*T*A*G*C*+A*+G*+C 3’*

*NHP MALAT1 ASO Sequence with LNA containing chemistry: 5’*

*+A*+G*+T*A*C*T*A*T*A*G*C*A*T*+C*+T*+G 3’*

### Antisense Oligonucleotide Synthesis

Oligonucleotides were synthesized using a MerMade 12 (LGC) DNA/RNA synthesizer on 100 µmol scale using standard solid-phase synthesis protocols. DNA and LNA phosphoramidites were purchased from Hongene Biotech Corporation. These include LNA: LNA-A(Bz), LNA-5MeC(Bz), LNA-T, LNA-G(dmf) and DNA: dA(Bz), dC(Ac), dT, dG(dmf). Constrained ethyl phosphoramidites (cET) were synthesized according to published procedures[61]. LNA-5MeC(Bz) solutions were prepared in a mixed solvent of DCM/acetonitrile (1:1, v/v). All other all other phosphoramidite solutions were prepared in acetonitrile in the presence of 3Å molecular sieves. 6-(Trifluoroacetylamino)-hexyl-(2-cyanoethyl)-(N,N-diisopropyl)-phosphoramidite was purchased from Glen Research (CAS: 133975-85-6; Glen Research Cat. # 10-1916). Each synthesis cycle was used to add one nucleotide and consisted of the following steps: detritylation, coupling, oxidation (or sulfurization), and capping. 5-Ethylthio-1H-tetrazole (ETT, Honeywell Research Chemicals, 0.25 M solution in acetonitrile) was used as an activator. Phenylacetyl disulphide (PADS, TCI Chemicals. A 0.2 M solution in 1:1 pyridine:acetonitrile was employed to introduce phosphorothioate linkages. Further details for solid-phase synthesis on 100 µM scale are shown in Table S5.

### Synthesis of 5’-hexylamino-modified ASOs

5’-hexylamino-modified ASOs were synthesized using the following solid phase oligonucleotide synthesis conditions: Using UnyLinker CPG support (500 Å), each coupling step used 4.1 eq amidite (0.05 M in MeCN) with 30 eq. ETT as activator (0.25 M in MeCN). The 5’-hexylamine spacer was introduced via 4.1 eq of 6-(Trifluoroacetylamino)-hexyl-(2-cyanoethyl)-(N,N-diisopropyl)-phosphoramidite (0.05 M in MeCN) with 30 eq. ETT as activator (0.25 M in MeCN). Oxidation steps used 0.02 M iodine solution in MeCN/pyridine/H_2_O (7:2:1 v/v/v) and sulfurization was achieved with 0.20 M PADS in pyridine/MeCN (1:1 v/v). After completion of solid-phase oligonucleotide synthesis, the 2-cyanoethyl protecting groups were cleaved using 20% diethylamine (DEA) in acetonitrile for 1 hour. Cleavage from solid support and nucleobase deprotection were achieved with NH_4_OH/EtOH (3:1) at 45°C for 20 hours. The resulting crude solution was concentrated under reduced pressure, and the solid residue was reconstituted in water for preparative reverse-phase HPLC purification using the conditions outlined in Table S6.

### Synthesis of 3’-amino modified ASOs

3’-amino-modified ASOs were synthesized using the same solid phase oligonucleotide synthesis conditions as described above but utilized 3’-PT-Amino-C6-CPG solid support (500A; 80-90 μmol/g, LGC group).

### Synthesis of 5’-maleimido modified ASOs

5’-maleimido-modified ASOs were prepared using the following procedure: To a solution of the purified 5’-amino-modified ASO (30 mg, by OD) in pH 6.0 PBS buffer (2.5 ml) was added a solution of 3-maleimidopropionic acid N-hydroxysuccinimide ester (MCOSu, 10 eq., TCI Chemicals) in DMF (2.5 ml). The resulting solution was shaken at room temperature for 18 hours after which the solution was desalted by passing through a sephadex G25 column (2.5×40 cm) on AKTA pure 25 M instrument, eluting with Milli Q water at 3 ml/min, monitored by UV 260/280 nm. The appropriate fractions were pooled and lyophilized.

### LC-MS Characterization data

5’-hexylamino ASO (mouse *Malat1*): calc. 5509.49, observed 5509.99

5’-hexylamino ASO (NHP *MALAT1*): calc 5510.48, observed 5511.17

3’-amino ASO (mouse *Malat1*): calc 5539.52, observed 5539.78

5’-maleimido ASO (mouse *Malat1*): calc 5660.61, observed 5660.93

5’-maleimido ASO (NHP *MALAT1*): calc for 5661.60, observed 5661.26

### Expression and purification of TV with S239C cysteine modification

The transport vehicle (TV) huIgG protein were expressed as knob-in-hole heterodimeric proteins via transient transfections of a CHO-K1 cell line. The cells were grown at 37°C, 5%CO_2_, 80% humidity, 150rpm and transfected using Mirus TransIT-PRO® transfection reagents according to the manufacturer’s instructions. Cultures were then temperature shifted to 32°C and harvested 7 days post transfection. The TV protein were then purified from the conditioned medium using Protein A chromatography (MabSelect PrismA, Cytiva, #17549802), followed by size exclusion chromatography using Superdex 200 (HiLoad 16/600 Superdex 200 pg, Cytiva, #28989335). Purified TV protein was buffer exchanged into PBS and stored at 4°C until further use.

### Bioconjugation, purification and analytics

The TV (10 mg/ml) generated from internal CHO cell line was first reduced using 30 molar equivalents of TCEP (Sigma, #646547) and 2 mM EDTA (Corning, #46-034-CI) 37°C for 1 hour. Reduction was confirmed by LC/MS (Ultimate 3000 LC, Orbitrap exactive EMR)). Post reduction, remaining TCEP was removed by dialysis using PBS, pH 6.8 containing 2mM EDTA. The TV was then re-oxidized with 50 molar equivalents of dHAA (Biosynth, #D-0300) at room temperature for 3 hours. Oxidation was confirmed by LC/MS. Next, 1.2 molar equivalents of linker-ASOs were added to the oxidized TV, and the mixture was incubated at room temperature for 1 hour. Unconjugated TV and conjugated TV with different OTR (Oligonucleotide to TV ratio) species were separated using anion exchange chromatography (Resource Q, Cytiva, #17117701) with mobile phase A: 50mM Tris, pH 7.5, and mobile phase B: 50 mM Tris, pH 7.5, 2M NaCl. OTR1 eluted between 30 - 40 % B was pooled and dialyzed into PBS. Final sample purity and OTR ratio was confirmed by reverse phase LC/MS (Mab pac RP, Thermo Fisher Scientific, #088644) with mobile phase A: 0.1 % formic acid in water, mobile phase B: 0.1 % formic acid in acetonitrile, and analytical size exclusion chromatography (Protein BEH SEC, Waters, #186005225) with mobile phase of 2x PBS. Final protein concentration was determined using the BCA assay (Thermo Fisher Scientific, #23225).

### Generation of an anti-ASO rabbit polyclonal antibody

Custom polyclonal antibody reagent campaign was done through collaboration with Rockland Immunochemicals, Inc (Pennsylvania, USA), following their standard immunization procedures. As previously described, a 3’amino-modified ASO against mouse *Malat1* was synthesized and directly conjugated to a KLH (keyhole limpet hemocyanin) peptide. Upon observation of desired titer response in male New Zealand rabbits and collection of production bleeds, the polyclonal was purified though a protein-A column.

### Characterization and validation of an anti-ASO pAb reagent

Initial specificity assessment of purified pAb was done as follows: standard ELISA (enzyme-linked immunoassay) plates were blocked with 3% Fish Gelatin in PBS (Rockland Immunochemicals, #MB-066-100). Antigen was coated to a final concentration of 5 μg/ml in TMBE (Rockland Immunochemicals, #TMBE-8000) substrate and treated with a concentration range of BSA (bovine serum albumin) as a protein control, BSA-conjugated immunogen sequence, immunogen sequence containing no phosphorothioate chemistry, and an unrelated sequence containing no phosphorothioate chemistry. Secondary antibody, Goat anti-Rabbit IgG (H&L) Peroxidase Conjugated (Rockland Immunochemicals, #611-1302) was diluted 1:10,000 in substrate. From this ELISA, the purified pAb was found to recognize the immunogen with a passing titer. No cross-reactivity was observed with BSA or sequences lacking phosphorothioate chemistry. To further characterize the selectivity and sensitivity of this reagent, the purified pAb was conjugated to act as a primary and secondary antibody in a generic MSD (Meso Scale-Discovery)-based sandwich electrochemiluminescence immunoassay. Primary antibody, biotinylated anti-ASO pAb, was added to blocked plates at a concentration of 1 μg/ml and a variety of ASOs were tittered and added to the plate at 1:20 dilution in 1% casein-PBS. Secondary antibody, ruthenylated (SULFO-TAG) anti-ASO pAb, was then added at a final concentration of 1 μg/ml. After appropriate washing, 1x MSD Read Buffer T (Meso Scale Discovery, Rockville MD) was then added to generate the electrochemiluminescence (ECL) signal. From this assessment, our rabbit anti-ASO reagent was found to be highly selective towards phosphorothioate containing ASO sequences, independent of sequence length and ribose chemistry. In the variety of ASOs assessed, an average sensitivity of down to 0.3 nM was observed in this immunoassay sandwich design.

### hTfR1 binding affinity with surface plasmon resonance (SPR)

Human TfR binding affinities of tested OTV variants were determined by SPR using a Biacore™ 8K instrument. OTV variants were immobilized on Cytiva Series S CM5 sensor chip (Cytiva, #29149603) using a Cytiva Human Fab capture kit (Cytiva, #28958325) at 10µg/mL at a flow rate of 10µL/min for 60 seconds. Serial 3-fold dilutions of recombinant human TfR apical domain at concentrations of 24.5, 74.0, 222, 667, and 2000 were injected at a flow rate of 30 μL/min for 60-seconds followed by a 300-second dissociation in a 1X HBS-EP+ running buffer (Cytiva, #BR100826). Following the injection, the chip was regenerated using 10mM glycine-HCl (pH 2.0) at the end of each cycle. Data analysis was conducted using Biacore Insight Evaluation software (version 2.0.15.12933). A 1:1 Langmuir kinetic binding model was used for hTfR binding affinity analyses.

### Fluorescence labeling of OTV for *in vitro* detection

Prior to labeling, 1 M Sodium bicarbonate (1/10 of sample volume) was added to the OTV (generation method above) to active lysine on the OTV. 1.2 molar equivalents of Alexa 647 were added to the OTV. The reaction mixture was incubated at room temperature for 1 hour. Desalting Zeba column equilibrated with 1x PBS pH 7.4 was used to remove excess Alexa 647. Concentration of labeled OTV was determined by BCA, and degree of AF 647 labeling was determined by A280 and A650.

### Primary hippocampal neuron culture and treatment

Primary neurons were prepared from the hippocampus of E16-E18 embryos of C57BL/6, CD-1 (Charles River Laboratories), or human transferrin receptor KI. Hippocampi were dissected in prechilled hibernate E solution. The cells were dissociated by ∼ 0.1 unit/hippocampus of papain at 37°C for 10 min and added DNAseI (3000 unit/ml) and FBS. After changing to NbActive 4 medium including Penicillin-Streptomycin, Glutamax, and 5-floro-2-deoxyuridine (final concentration 1 µM), the cells were dissociated by gently pipetting and plating on PDL-coated 96 well plates at the density 25K per well. For RT-qPCR assay, ASO, TV or OTV were added at concentrations ranging from 12 – 500 nM 3 days after seeding and incubated for 7 days. For cellular uptake assay, ASO, TV or OTV were added at 500 nM 7 days after seeding, and cells were fixed at specific time points.

### *Malat1* RNA level in fibroblast cells or Neuro-2a cells

Cynomolgus monkey brain vascular fibroblasts (Cell Biologics, #MK-6076) cells or Neuro-2a cells were seeded on PDL-coated 96-plate. Malat1 ASO was treated at 40nM with lipofectamine 3000 one day after plating cells. Fibroblast cells were incubated for 5 days and performed RT-qPCR. Neuro-2a cells were incubated for 3 days and performed RT-qPCR.

### ASO Immunocytochemistry

Cynomolgus monkey brain vascular fibroblasts cells or SH-SY5Y cells were seeded on PDL-coated 96-plate. After treating ASOs and incubating for 3-5 days, cells were fixed with 4% PFA in PBS at room temperature. Cells were treated with blocking buffer (0.05% Tween20 and 1% BSA in PBS) for 1 hour and then incubated with anti-phosphorothioate polyclonal antibody (pAb, Rockland, 0.66 µg/ml) in blocking buffer at 4°C overnight. After three washes using PBS, cells were incubated with fluorescently labeled secondary antibodies with CellMask (Thermo Fisher Scientific, #H32721) in a blocking buffer at room temperature for 1 hour. Immunostained cells were imaged using the Opera Phenix High Content Screening System (PerkinElmer). Acquired images were analyzed by using the Harmony software (PerkinElmer).

### ASO and OTV cellular uptake assay

SH-SY5Y cells were seeded 4K per well in PDL coated 96 well plates in 100 µl of complete growth medium and incubated overnight at 37°C in 5% CO_2_ incubator. To label organelles, 2μl/well of CellLight, BacMam2.0 reagent (Thermo Fisher Scientific) corresponding to each organelle was added and incubated overnight. Each labeled ASOs, ATV or OTVs were diluted 40 nM in Opti-Mem before being administered to cells. Live cell imaging was performed at 37°C in 5% CO_2_ for 24 hours using the Opera Phenix High Content Screening System (PerkinElmer). Acquired images were analyzed by using the Harmony software (PerkinElmer).

### Super resolution confocal imaging

SH-SY5Y cells were plated on glass bottom 96 well plates (Greiner Bio-One, #655891) after coating Poly-L-Lysine for 15 min at room temperature. 24 hours after 500 nM ASOs or OTVs treatment, cells were fixed with 4% PFA in PBS for 30 min. Cells were treated with blocking buffer (0.05% Tween20 and 1% BSA in PBS) for 1 hour and then incubated with primary antibodies in blocking buffer at 4°C overnight. After three washes using PBS, cells were incubated with fluorescently labeled secondary antibodies in a blocking buffer at room temperature for 1 hour. ages were captured using a laser scanning confocal microscope (Leica SP8; Leica Microsystems, Inc). It operated in super-resolution LIGHTNING mode acquired with a 63x/1.4 NA oil objective at a pixel size of 40 nm and processed using an adaptive processing algorithm. Format size was 2480x2480 with a speed 8000, Image sequential scanned using frame mode with 10 frame average. The representative image was made by three-dimensional reconstruction using Imaris (Bitplane). By using surface rendering, and fluorescence thresholding, we were able to obtain an accurate 3D structure.

### huIgG ELISA protocol

Total human IgG concentrations in murine plasma and tissue homogenates were quantified using a generic anti-human IgG sandwich-format (ELISA). Briefly, plates were coated overnight at 4°C with donkey anti-human IgG pAb (Jackson ImmunoResearch, #709-006-098) at 1 µg/ml in sodium bicarbonate solution (Sigma, #C3041-50CAP). Following incubation and plate wash with buffer (PBS + 0.05% Tween20), prepared test samples (with sample pre-dilution, where appropriate, in PBS + 0.05% Tween20 + BSA (10 mg/ml)) and relevant standards were added to the assay plate and were allowed to incubate for 2 hours. Following test sample incubation and wash step, secondary antibody, goat anti-human IgG (Jackson ImmunoResearch, #109-036-098), was diluted in blocking buffer (PBS +0.05% Tween20 + 5% BSA (50 mg/ml)) to a final concentration of 0.02 µg/ml. Following a 1 hour incubation and final wash step, plates were developed by adding TMB substrate (Thermo Fisher Scientific, #34028) and incubated for 5-10 minutes. Reaction was quenched by adding 4N H_2_SO_4_ (Life Technologies, #SS03) and read using 450 nM absorbance. Total human IgG concentrations in cynomolgus plasma and tissue samples were quantified using a generic MSD-based sandwich electrochemiluminescence immunoassay. In brief, MSD GOLD 96-well small-spot streptavidin-coated microtiter plates (Meso Scale Discovery, Rockville, MD) were treated with a blocking step of 1% casein-based PBS (Thermo Scientific, Waltham, MA) for approximately 1 hour. Following a wash step, a biotinylated goat anti-human IgG polyclonal antibody (SouthernBiotech, Birmingham, AL) diluted at a working concentration in 1% casein-PBS was added to the plate and incubated for 1-2 hours. Following plate coating and wash steps, test samples (standards, QCs, and unknowns) were added to the assay plate and allowed to incubate for 1-2 hours. Following the sample incubation and subsequent wash step, a monkey plasma pre-adsorbed secondary ruthenylated (SULFO-TAG) goat anti-human IgG polyclonal antibody (Meso Scale Discovery, Rockville, MD) at a working concentration in 1% casein-PBS was added and incubated for 1 hour. A 1x MSD Read Buffer T (Meso Scale Discovery, Rockville MD) was then added to generate the electrochemiluminescence (ECL) signal. For the above two species specific methods, all assay reaction steps were performed at ambient temperature with gentle agitation (where appropriate). All test samples were pre-diluted at the assay minimum-required-dilution (MRD) of 1:100 prior to analysis. The assay standard curves were fitted with a weighted four-parameter (4P) nonlinear logistic regression for use in calculating concentrations of unknown samples.

### Intact OTV ECLIA protocol

Quantification of Intact OTV in plasma and tissue homogenates were measured using a method similar to a modified hybridization-based electrochemiluminescence immunoassay (ECLIA)[62]. Briefly, custom biotinylated capture probe (Integrated DNA Technologies, Custom) at working concentration described was combined with prepared test samples (with sample pre-dilution, where appropriate) and relevant standards in TE buffer (10 mM Tris-HCl containing 1mM EDTA). Prepared samples in TE buffer were added, in a 1:1 mix, into 2x SSC Buffer (Sigma-Aldrich, #AM9770) containing hybridization capture probe at 25 nM. Samples were hybridized at 50°C for 45 mins. Following the incubation, hybridized OTV-probe products were added to the wells of an MSD GOLD 96-well streptavidin-coated microtiter plate (Meso Scale Discovery, Cat#L15SA) and incubated for approximately 30 mins. Following incubation and plate wash step, secondary ruthenylated (SULFO-TAG) goat anti-human IgG antibody (Meso Scale Discovery, #R32AJ-1) at a working concentration of 0.5 µg/ml in assay diluent (SuperBlock™ T20 (Thermo Fisher Scientific, #37536)) was added to the assay plate and incubated for approximately 1 hour. Following a plate wash, MSD Read Buffer T (Meso Scale Discovery, #R92TC) was then added to generate the electrochemiluminescence (ECL) assay signal. All assay plate steps were performed at ambient temperature with gentle agitation on a plate shaker (where appropriate); and all test samples were pre-diluted at the assay MRD of 1:20 prior to analysis. Sample ECL signals generated in the assay were subsequently processed into concentrations by back-calculating off the assay calibration (CS) curve. The assay CS curve was fitted with a weighted 4PL nonlinear logistic regression to calculate the concentration of unknown samples. Sequence for capture antisense probe is described below where ‘/5BioTEG/’ denotes a 5’-biotin with triethylene glycol (TEG) linker and ‘+’ denotes an LNA base modification to the subsequent base.

*Mouse Capture probe: /5BioTEG/G CT+G +CTA TTA +GAA TGC*

*NHP Capture probe: /5BioTEG/CA+GAT+G+CTATA+GTACT*

### Total ASO ECLIA protocol

Quantification of total ASO (in OTV conjugated and liberated forms) in plasma and tissue homogenates were measured using a method similar to a modified hybridization-based ECLIA assay[63]. Briefly, custom capture and detection probes (Integrated DNA Technologies, Custom) at working concentrations described were combined with prepared test samples (with sample pre-dilution, where appropriate) and relevant standards in TE buffer. Prepared samples in TE buffer were added, in a 1:1 mix, into 2x SSC Buffer (Sigma-Aldrich, #AM9770) containing hybridization probes at 25 nM and recombinant proteinase K enzyme (Thermo Fisher Scientific, #28352) at 100 µg/ml. Hybridization/Enzyme mixture was then digested, denatured, annealed, and cooled in the following manner in a thermalcycler: 50°C for 15 mins, 95°C for 5 mins, 40°C for 30 mins, 12°C for ∞. Following hybrid product formation, samples were added to the wells of an MSD GOLD 96-well streptavidin-coated microtiter plate and incubated for approximately 30 mins. Following incubation and plate wash step, secondary SULFO-Tagged sheep anti-digoxygenin antibody (Novus Biologicals, #NB100-65495) at a working concentration of 0.5 µg/ml in assay diluent and incubated for approximately 30 mins. Following a final plate wash, MSD Read Buffer T was added to generate the plate ECL signal. All assay plate steps were performed at ambient temperature with gentle agitation on a plate shaker (where appropriate); and all test samples were pre-diluted at the assay MRD of 1:20 prior to analysis. Sample ECL signals generated in the assay were subsequently processed into concentrations by back-calculating off the assay CS curve. The assay CS curve was then fitted with a weighted 4PL nonlinear logistic regression to calculate the concentration of unknown samples. Sequences for capture and detection probes are described below where ‘/3Dig_N/’ denotes a 3’-amide linked digoxygenin.

*Mouse Capture probe: /5BioTEG/G+C+T+G+CTAT*

*Mouse Detection probe: T+A+GAA+T+GC/3Dig_N/*

*NHP Capture probe: /5BioTEG/CA+G+AT+GC NHP*

*Detection probe: T+ATA+G+TA+CT/3Dig_N/*

### Pharmacokinetic calculations

Analysis of pharmacokinetic trends, through a non-compartmental approach, was done using Dotmatics Software 5.5 (Boston, Mass.). Parameters were estimated using nominal sampling times relative to administration start. Any samples that were below detection limits (BOD) were omitted. To calculate exposures, a linear up & log down model, was used to calculate pharmacokinetic exposures. Descriptive statistics were also generated using Dotmatics.

### Mouse handling and tissue collection

Animals received care in accordance with the Guide for the Care of Use of Laboratory Animals, 8^th^ Edition. Animals were house in groups, not exceeding 5 animals per cage. Mouse husbandry and experimental procedures were approved by the Denali Institutional Animal Care and Use Committee. C57BL/6 and TfR^mu/hu^ KI mice [23] were acquired from The Jackson Laboratory (Sacramento, CA). Housing conditions included standard pellet food and water provided ad libitum, a 12-hour light/dark cycle at a temperature of 22°C with cage replacement once every three weeks and regular health monitoring. Mice were peripherally administered therapeutic treatment via intravenous (IV) tail vein injection (∼150 µl total volume). For tissue collection, animals were anesthetized with 2.5% tribromoethanol and whole blood collected via cardiac puncture into EDTA coated tubes for plasma drug concentration assessment. Following transfer to EDTA coated tubes, whole blood was spun down at 12,700 rpm for 7 min at 4°C before collecting the top plasma layer. Mice were then perfused with ice-cold PBS transcardially at a rate of 5 ml/min for 3-5 min. For biochemical analysis, tissues were collected, weighed, snap frozen on dry ice, and then stored at -80°C. For immunohistochemistry analysis, tissues were collected and subsequently drop fixed in 4% paraformaldehyde (PFA) for 24-48 hours at 4°C and then transferred to PBS + 0.1% sodium azide until ready for sectioning. Detailed mouse information for all in vivo studies in the manuscript can be found in Table S7.

### Intracerebroventricular Bolus (ICV) surgery

Procedure was slightly adapted from a previous publication[64]. In brief, the surgical area was sterilized with isopropanol and surgical instruments were autoclaved. Mice were brought under anesthesia with ∼4% isoflurane. Hair was shaved from the shoulder region to between the eyes prior to placement on the stereotax surface. 20uL of 0.5% Bupivicain was administered subcutaneously to numb the area prior to incision. With a maintenance level of ∼2% isoflurane, an incision was made from the base of the neck to between the eyes. Following cleaning with hydrogen peroxide, the skull was punctured with a drill bit and the injection needle was slowly inserted into the brain at the following coordinates: ML +1, AP +0.3, DV -3 (from brain surface). After waiting one minute to allow for brain sealing around the needle, a dose of 10 µl ASO diluted in saline was administered at 1 µl per second. Two minutes later, the needle was raised slowly out of the brain. Following ICV bolus, the incision was sutured and mice were treated with 0.2mL of 0.5 mg/mL Carprofen for pain. Mice were then transferred to a cage placed over a heated recovery pad and observed for full recovery. Mice were monitored daily after surgery to check for pain, discomfort, or infections.

### Non-Human Primate handling and tissue collection

Female cynomolgus monkeys (Mainland Asian origin) aged 2-4 years were housed in groups of 2, 3, or 4 animals (except for procedures that require individual housing) in appropriately sized caging. Housing and handling follow specifications as outlined in the USDA Animal Welfare Act (9 CFR, Parts 1, 2, and 3) and as described in the *Guide for the Care and Use of Laboratory Animals* in an AAALAC, International accredited facility. All procedures were conducted following the Test facility’s Standard Operating Procedures (SOP). IV dosing of test articles were administered via the saphenous vein with a slow bolus injection of approximately 1 min – 1.1mg/kg for naked ASO^Mf^ and 30mg/kg for OTV. IT drug administration (4mg naked ASO^Mf^) was performed with animals under anesthesia with animals placed in the left lateral recumbency position[65]. Using a 22-gauge Gerti Marx spinal needle, the lumbar cistern was percutaneously punctured at the approximate L4/L5 region, using aseptic technique. Positive CSF flow was confirmed through the needle for correct needle placement. Test article syringe was then attached, and the test article was slowly infused by hand as a slow bolus over approximately 1 mL/minute for a total volume of 2mL. After dosing, brief pressure was applied to the site of injection, anesthesia was reversed, and the animals were allowed to recover naturally. In-life blood sampling for plasma pharmacokinetics was collected from the femoral vein/artery. Collections were transferred to EDTA coated tubes, whole blood was spun down at 12,700 rpm for 7 min at 4°C before collecting the top plasma layer. For terminal collections, animals were euthanized and perfused with cold PBS containing heparin (1000 U/L) via the left ventricle using a roller pump for at least 5 mins. For Brain, left hemisphere was used for frozen tissue punches and the right hemisphere was used for fixation. Brain along with cervical, thoracic, and lumbar spinal cord samples were collected and sliced coronally into 4mm thick slabs[66]. For biochemical analysis, triplicate samples of all detailed individual brain regions were collected from the slabs using 3 or 2mm punches with care taken to avoid at least 2mm minimum distance from brain surfaces that may contact CSF in the ventricles or subarachnoid space for all regions except the frontal cortex surface punches. Samples were collected, weighed, and snap frozen on dry ice, and stored at -80°C until analysis. Samples collected for immunohistochemistry (IHC) were preserved in 4% paraformaldehyde (PFA). Detailed NHP study animal and design details can be found in Table S8.

### Tissue homogenization

Weighed frozen tissue samples were processed for biochemical assays by adding 5X or 10X volume of chilled 1% NP40 + PBS homogenization buffer with added cOmplete Protease Inhibitor (Roche #04693132001) and PhosStop (Roche 04906837001) phosphatase inhibitors. Samples were homogenized using 3 mm tungsten carbide beads in 1.5 ml Eppendorf tubes and shaken using the Qiagen TIssueLyzer II (Cat No./ID: 85300) for 6 min at 27 Hz. For assays requiring purified tissue lysate (e.g., Western blot), samples were centrifuged at 14,000 x g for 15 min at 4°C and a BCA assay was subsequently performed for total protein normalization. For total ASO quantification, crude homogenate was used for analyses.

### Western blot

Tissue lysates, as prepared above, were protein-normalized using a BCA assay (Thermo Fisher Scientific, #23225). Protein normalized lysates were mixed with loading buffer (Thermo Fisher Scientific, #NP0007) and reducing agent (Thermo Fisher Scientifc, #NP0004) to a final protein concentration of approximately 10 mg/mL. 10 uL of each sample was resolved by electrophoresis using a 4-12% NuPAGE (Thermo Fisher Scientific, #NP0335BOX). Semi-dry transfer was performed using the Trans-Blot Turbo System (Bio-Rad, #1704150) with 0.2-µm PVDF transfer packs (Bio-Rad, #17001917). Transferred blots were blocked with 5% milk in TBS-T containing 0.05% Tween20 and incubated with primary antibody overnight. LICOR secondary antibodies were used for detection. Washes were performed using TBS-T. Blots were imaged by the Li-COR Odyssey imaging system. Band signal intensity was using FIJI software.

### RNA extraction protocol

*In vitro samples*: RNA isolation from primary hippocampal neurons using RNeasy plus micro kit (Qiagen, #74034), according to the manufacturer’s recommendations.

*In vivo samples*: One ml of cold TRIzol^TM^ Reagent (Invitrogen, #15596026) was added to each tissue sample and homogenized using a TIssueLyzer II (Qiagen, #85300) with 3 mm stainless steel beads at 27 Hz for 6 minutes. 0.2 ml of chloroform was then added to lysed samples and allowed to incubate at ambient temperature for 2-3 mins. Samples were then centrifuged for 15 mins at 12,000 x g at 4°C. While kept on ice, the aqueous phase containing RNA was transferred to a new tube. Isopropanol at a volume of 0.5 ml per 1 ml of TRIzol^TM^ was added, vortexed vigorously, and incubated for 10 mins on ice. Following incubation, samples were centrifuged for 10 mins at 12,000x g at 4°C to form RNA precipitate. Upon removal of supernatant, 0.5 ml of 75% ethanol per 1 ml of TRIzol^TM^ was added to wash and samples were centrifuged for 5 minutes at 7,500 x g at 4°C. Supernatant was aspirated and RNA pellets were resuspended in 100 µl of RNase-free water (Ambion, #AM9937).

### Real time quantitative PCR (RT-qPCR) assay and data analysis

To evaluate target RNA levels, RT-qPCR was performed using either 75ng of RNA extracted from *in vitro* cell lysates or 40 ng RNA extracted from bulk mouse tissues. For *in vitro* samples, cDNA was generated with the SuperScriptTM IV VILO Master Mix (Thermo Fisher, #11756050) by incubating the reaction at 25°C for 10 min, followed by 50°C for 10 min and 85°C for 5 min. The Taqman Fast Advanced Master Mix (Thermo Fisher, #4444557) was used for the subsequent RT-qPCR protocol. For *in vivo* samples, target RNA levels were evaluated using the Express One-Step kit (Thermo Fisher, #1178101K). All experiments used the following Taqman probes: MmMalat1 (Thermo Fisher, #Mm01227912) and MmGapdh (Thermo Fisher, #Mm99999915) for cell lysates and mouse tissue; MfMALAT1 (Thermo Fisher, #10336022) for cynomolgus monkey tissue. For *in vitro* samples and *in vivo* mouse tissues, *Malat1* RNA levels were normalized to the housekeeping gene *Gapdh*. Cynomolgus monkey tissues were normalized to total RNA input due to variability in *GAPDH* levels between monkeys. RT-qPCR was performed by running the following thermocycler program on a QuantStudio 6 Flex system (Applied Biosystems): 50C for 15 min, 95C for 2 min, then 40 cycles of 95C for 15 sec, 60C for 1 min. Average CT values were measured for each gene from technical duplicates. After normalizing to either *Gapdh* or total RNA input, delta CT values were calculated relative to the control group and plotted as relative expression levels.

### Brain immunofluorescence imaging

Following post-fixation in 4% PFA, mouse hemibrains were coronally sectioned at a thickness of 40 µm. Floating sections were then incubated at room temperature for 2 hours while rocking in blocking buffer (PBS + 5% BSA + 5% NDS + 0.5% triton X-100) then transferred to antibody dilution buffer (blocking buffer plus primary antibodies; see reagents table). Sections were incubated for 48 hours at 4°C on a rocker. After primary antibody incubation, sections were washed 3X in PBS and transferred to antibody dilution buffer (blocking buffer plus secondary antibodies at 1:1000) and incubated at room temperature for 2 hours on a rocker. Samples were subsequently washed with PBS+DAPI (Invitrogen, #D1306 1:10,000) for 15 min, then washed 3X with PBS before mounting on 2” x 3” microscope slides with Prolong Glass hardset mounting media (Life Technologies, #P36984). Slides were allowed to dry overnight at room temperature before imaging.

Image acquisition, processing, and quantification:

Immunostained sections were imaged with a Zeiss AxioScan slide scanner running Zen 3.5 software. Each tissue section was entirely imaged by scanning across the identified tissue area, taking images of each location with exposure settings and filter sets to detect DAPI and Alexa-488, Alexa-555, and Alexa-647-tagged antibodies. Automated shading correction and stitching in Zen was then used to produce single complete images for each section. Identical imaging settings and post-processing methods were used to collect images from all tissue sections from a given experimental run.

To quantify ASO localization and distribution across tissue sections, images were first processed to identify specific classes of objects using sequences of advanced image processing tools in custom macros designed in-house with Zeiss Zen 3.5 software. ASO, NeuN (neurons), and DAPI (nuclei) stained objects were detected by local thresholding and size selection to remove too-small staining artifacts. Colocalization objects were then defined as areas of overlap between NeuN and ASO objects, DAPI and ASO objects, NeuN-negative areas and DAPI (non-NeuN nuclei), and ASO overlap with non-NeuN nuclei. Total tissue area was identified by standard thresholding of heavily smoothed images in the non-specifically stained Alexa 488 channel (high tissue background staining). Only ASO, NeuN, DAPI, and overlap objects located within total tissue area were included in subsequent quantifications. To assess broad patterns of ASO distribution, the Alexa-647 channels of images were heavily smoothed (Median 249) and included as a separate channel for quantification in subsequent steps.

Once classes of objects were identified, we quantified the total count, total area, and total sum intensity of the Alexa-647 (ASO) channel for the total tissue area and each set of objects in every imaged tissue section. We also quantified the standard deviation of staining intensity levels across the total tissue area of both the standard and heavily-smoothed Alexa-647 channels to assess whether distribution of ASOs were smooth or locally concentrated across the tissue.

### Nuclei isolation, capture, and single nuclei library preparation (Fig. 3 snRNA-seq)

Sixteen mouse brain tissue samples were divided into four batches for sample processing, each batch containing one replicate sample from all four experimental groups. Nuclei were isolated with a method similar to a previously published protocol [67] using Nuclei EZ Prep buffer (Sigma Aldrich, # NUC101) containing a final concentration of 1X protease inhibitor (Complete, EDTA-free (Roche, #11873580001) and 0.4 U/µl RNase inhibitor (Ambion, #AM2682). For each sample, approximately 80 mg of tissue was dissociated with 25 strokes each of pestle A and B using a 2 ml glass dounce homogenizer and the lysates were incubated on ice for 15 minutes. Afterward, the suspension was centrifuged at 500g in 4°C for 5 minutes, the pellet was resuspended in Nuclei EZ Prep buffer and incubated for another 15 minutes on ice.

After lysis, the nuclei were centrifuged at 300g in 4°C for 5 minutes and the pellet was resuspended in 500 µl of 1% BSA in PBS. Purification was performed using a 2 M sucrose cushion with centrifugation at 13,000g in 4℃ for 45 minutes. The pellet was resuspended in 2% BSA in PBS and passed through a 30-µm strainer. Concentrations of nuclei were determined using a Vi-CELL cell counter (Beckman Coulter).

Single nuclei were captured on a 10x Genomics Chromium Controller with the Chromium Next GEM Single Cell 3’ kit (version 3.1) targeting a recovery of 10,000 nuclei per sample. Next-generation sequencing library preparation was performed as per manufacturer’s protocol CG000204 (10x Genomics, Revision B). One sample failed to be captured, reducing the total number of samples to 15. The number of captured nuclei in each library was estimated based on a shallow sequencing run on an Illumina MiSeq instrument. Libraries were pooled in proportions that ensured approximately similar sequencing coverage per cell across samples and were sequenced on an Illumina NovaSeq 6000 instrument by Seqmatic (Fremont, CA). For each sample, a minimum of 200 million reads pairs were obtained.

### Nuclei isolation, capture, and single nuclei library preparation (Fig. 4 and Fig. 6 snRNA-seq)

Eighteen mouse brain tissue samples were divided into four batches for sample preparation and single nuclei capture, each batch containing at least one sample per experimental group. Nuclei were isolated from approximately 80-100 mg of frozen mouse brain tissue samples in a procedure similar to a previously published “Frankenstein” protocol (https://www.protocols.io/view/frankenstein-protocol-for-nuclei-isolation-from-f-bq23mygn).

Briefly, frozen tissue samples were placed into chilled Nuclei EZ Lysis Buffer (Sigma Aldrich, # NUC101) and mechanically homogenized using a 2 ml dounce homogenizer by stroking 15-20 times with pestle A followed by 15-20 strokes of pestle B. The lysate was incubated on ice for 5-10 mins before filtering through a 70 µm strainer mesh, and then pellet and resuspended again in Nuclei EZ Lysis Buffer for another 5 mins of incubation on ice. After 3-4 washes to remove debris at 500g in 4°C for 5 minutes, nuclei were resuspended in Nuclei Wash Buffer with a final concentration of 0.2 U/µl RNase inhibitor (Ambion, cat. no. AM2682) and 1% BSA in PBS. The nuclei solution was stained with DAPI at a final concentration of 10 µg/ml and then filtered through a 30 µm strainer. Roughly 25,000 - 35,000 nuclei were FACS sorted on BD FACSAria (Becton Dickinson) directly into 10x Genomics RT master mix for a target recovery of 10,000-15,000 nuclei. RT enzyme was added immediately prior to microfluidic loading onto the Chromium controller for nuclei capture, and libraries were prepared per 10X Genomics user guide (CG000204 Rev B).

A shallow sequencing run was performed on an Illumina MiSeq instrument to estimate the number of nuclei captured per library. This capture information was used to pool libraries in equal proportions for sequencing on the Illumina NovaSeq 6000 instrument generating paired-end reads (28x10x10x90 bases) performed by SeqMatic (Fremont, CA).

### Nuclei isolation capture, and single nuclei library preparation of non-human primate samples

Nine 20 mg thalamus tissue samples from non-human primates were divided into two batches of four and five samples respectively, each batch covering all three experimental groups. Dissociation was performed using a protocol similar to the Frankenstein protocol mentioned above.

### Processing of Figure 3 and Figure S2 snRNA-seq data

Filtered counts matrices generated with Cell Ranger v6.1.1 were further processed in R v4.1.3 using Seurat v4 [68]. Nuclei with >=50 genes and <=5000 genes detected were retained. Individual Seurat objects were merged. Data were normalized using the NormalizeData, highly variable genes were detected using the VST method in the FindVariableFeatures, and the data were then scaled using the ScaleData function. Dimensionality reduction was performed in Seurat with default settings.

### Processing Figure 4 and Figure S3 snRNA-seq data

Filtered counts matrices generated with Cell Ranger v6.1.1 were further processed in R v4.1.3 using Seurat v4. Nuclei with >=250 genes and <=4500 genes detected were retained. Individual Seurat objects were merged, excluding one mouse that showed poor perfusion. Data were normalized, and highly variable genes were identified. Data were then scaled with gene detection depth regressed out. Dimensionality reduction was performed in Seurat with default settings.

### Doublet Detection

Probable doublet nuclei were identified using the scDblFinder package [69]. Cluster labels were provided to scDblFinder by performing unbiased clustering in Seurat with a resolution of 0.1. Doublet detection was then carried out using the scDblFinder function with default parameters. Doublets were labeled and then excluded from downstream analyses.

### Predictive cell type labeling

Cell type labels were assigned using the label transfer functionality in Seurat. The Mouse Brain Atlas (MBA) single-cell data set [38] was used as a reference for label transfer. This reference data and our snRNA-seq data were anchored by CCA, and cell type labels were then predicted and transferred, using the FindTransferAnchors and TransferData functions in Seurat, respectively. For the whole brain cell type labeling, nuclei that could not be confidently assigned granular taxonomy labels (e.g.: astrocytes, oligodendrocytes, vascular endothelium, pericytes, etc.) were left with broad taxonomy labels (e.g.: glia and vascular cells). Granular neuron subtyping was then performed by iterative subclustering and label transfer using the neuronal subtypes in the MBA.

### Pseudobulk differential expression testing

Malat1 expression differences in snRNA-seq data were assessed by differential expression testing in pseudobulk data. Pseudobulk counts were generated by summing counts across cell types within biological replicates using the aggregate AcrossCells function in the scuttle package. Differential expression analysis was then carried out using the standard DESeq2 pipeline[39]: DESeq was run with default parameters (Wald test for significance) using a model design of “∼ treatment + batch” across each cell type.

### Pseudospace analysis of *Malat1* knockdown across cortical depth

Cortical depth was predicted using the pseudotime functionality in Monocle 3[70] driven by genes with distinct cortical expression patterns. Cortical neurons were identified by label transfer and subclustered. The expression matrix was then subsetted to include only genes with >70% probability of enrichment in one or more cortical layers as predicted by Belgard et al, 2011. UMAP dimensionality reduction was carried out in Seurat using only these genes. The resulting UMAP was then used to calculate pseudotime in Monocle 3. Scaled expression values were then plotted against pseudotime using ggplot2, and a local regression line was fit to represent the moving average across pseudospace in each treatment group.

### Mapping of Cynomolgus Monkey raw reads and custom reference generation

We created a custom *Macaca fascicularis* genome index for use with Cellranger (10X Genomics). Genome assembly 6.0 (GCA_011100615.1) and matching gene annotations were downloaded from Ensembl release 106. The annotation of non-coding RNAs such as MALAT1 in the *Macaca fascicularis* genome is incomplete. Ensembl (release 106) annotates the human MALAT1 gene at position 11:65,498,546-65,507,374 (GRCh38.p13). This 20 kb region aligns to position 14:8,898,660-8,903,763 in the *Macaca fascicularis* genome, which contains the non-coding ENSMFAG00000062016 gene, the *Macaca fascicularis* MALAT1 ortholog.

We observed that alignments from both bulk whole-transcriptome and 3’ single-nuclei RNA-seq *Macaca fascicularis* data extend beyond both ENSMFAG00000062016’s 5’ and 3’ ends, indicating the gene is longer than currently annotated. To capture all RNA-seq reads originating from this locus, we expanded the *Macaca fascicularis* MALAT1 gene region to coordinates 14:8,898,273-8,905,739.

The custom Cell Ranger index was created following the instructions provided by 10X Genomics. First, gene annotations were filtered to retain the following gene-biotypes: protein_coding, lncRNA, antisense, IG_LV_gene, IG_V_gene, IG_V_pseudogene, IG_D_gene, IG_J_gene, IG_J_pseudogene, IG_C_gene, IG_C_pseudogene, TR_V_gene, TR_V_pseudogene, TR_D_gene, TR_J_gene, TR_J_pseudogene, TR_C_gene. Afterward, the genome index was created with the cellranger mkref command with default options. Reads were subsequently mapped using this new reference with Cell Ranger v7.0.1.

### Processing *macaca fascicularis* snRNA-seq data for Figure 6 and Figure S8

Filtered counts matrices generated with Cell Ranger were further processed in R using Seurat v4. Nuclei with >=800 and <=6000 genes detected were retained. In addition, nuclei with >30,000 UMIs and >1% ribosomal reads were removed. To account for apparent technical noise in broad gene expression, individual sample gene expression matrices were combined by the Seurat integration functionality, utilizing CCA. This was performed using the FindIntegrationAnchors and IntegrateData functions in Seurat v4, using 2000 shared variable features. Integrated expression values were then scaled and PCA was performed. Dimensionality reduction and clustering were performed in Seurat with default settings. Doublet detection was performed using scDblFinder as before with the mouse snRNA-seq data sets. However, after automated doublet removal, a few remaining apparent doublet clusters were retained. These clusters were identified by marker gene expression and removed from further analysis.

### snRNAseq datasets

All raw snRNA-seq data for both mouse studies and the cynomolgus study were uploaded to NCBI Gene Expression Omnibus (GEO) and can be found using the following accession numbers: GSE201741, GSE201787, and GSE208114.

### Capillary depletion

Capillary depletion was performed as previously described[23]. Immediately post PBS perfusion, hemi-brains with the meninges, choroid plexus, and cerebellum all carefully removed were collected into ice chilled HBSS buffer. Brains were homogenized using 10 strokes with a dounce homogenizer that used a smaller diameter pestle. The resulting homogenate was spun at 1,000g for 10 min at 4°C and the supernatant discarded. The pellet was then carefully resuspended in 2 ml of 17% dextran (MW 60,000; Sigma, #31397) and centrifuged at 4,122g for 15 min at 4°C. The resulting pellet was saved as the vasculature fraction. The supernatant containing brain parenchymal cells and myelin was transferred to a new tube, diluted with HBSS to wash the cells, and then centrifuged at 4,122g for 15 min at 4°C. The resulting pellet contained vascular-depleted parenchymal cells. Following the isolation, both vascular and parenchymal pellets were resuspended in cold 1% NP40 in PBS buffer with protease and phosphatase inhibitors added (Roche 04693159001 and 04906837001). The resuspended pellets were then vortexed for 20 s, incubated at 4°C for 20 min, and centrifuged at 12,700g for 10 min at 4°C. Lysate total protein concentration was measured using BCA assay (Thermo Fisher Scientific, #23225)

### Statistical Analysis

All non-sequencing data was analyzed using either a Student’s t-test, one way analysis of variance (ANOVA), or two-way ANOVA with multiple comparisons when indicated. Data were plotted and statistical tests were performed using GraphPad Prism. Half-lives were calculated in GraphPad Prism using a non-linear regression model. Detailed statistical analyses for sequencing data is described in the snRNAseq section above. See Tables S7-8 for animal numbers included in all *in vivo* studies.

## Acknowledgments

The authors would like to acknowledge collaborators Kaitlyn Whalen, Georgia Sfyroera, and Dan Kenney from Rockland Immunochemicals, Inc. for their contributions to the generation of the anti-ASO rabbit polyclonal antibody. The authors would like to thank Do Jin Kim and Yaneth Robles Colmenares for their input on protein binding experiments. The authors would like to thank Hilda Solanoy, Amy Leung, Roni Chau, Butch Benitez, Isabel Becerra, Kevin Rebadulla, Hoang Nguyen, and Meredith Calvert for their contributions towards *in vivo* study coordination, troubleshooting, and support.

## Funding

Funding for this work was provided by Denali Therapeutics, Inc.

## Author contributions

Conceptualization: SJB, MBT, CK, DT, MS, RJW, LK, MSD, JWL, SLD

Methodology: SJB, MBT, CK, DT, MS, RD, LN, RCW, YZ, KSC, CBD, JD, CH, MP, AS, CMMK, RT, TS, LK, SLD

Investigation: SJB, MBT, CK, DT, MS, RD, LN, RCW, YZ, MA, AC, KSC, JC, AC, CBD, JD, CD, TE, CH, DH, MP, ER, AS, RT, HT, TS, SLD

Visualization: SJB, MBT, CK, DT, MS, LK, SLD Funding acquisition: -

Project administration: SJ, WK, LS

Supervision: RCW, CM, JZ, AAE, KG, CMMK, PES, RGT, RJW, TS, LK, MSD, JWL, SLD

Writing – original draft: SJB, MBT, CK, DT, MS, LK, SLD

Writing – review & editing: SJB, MBT, CK, DT, MS, LN, RCW, YZ, MA, JD, CH, SJ, MP, AS, RT, HT, RGT, RJW, LK, FR, MSD, JWL, SLD

## Competing interests

All authors, except MA, AC, CD, and FR, are full-time employees and/or shareholders of Denali Therapeutics, Inc. MA, AC, CD, and FR are full-time employee and/or shareholder of Ionis Pharmaceuticals, Inc. This work has been in part described in one or more pending patent applications.

---

**Fig. S1.**
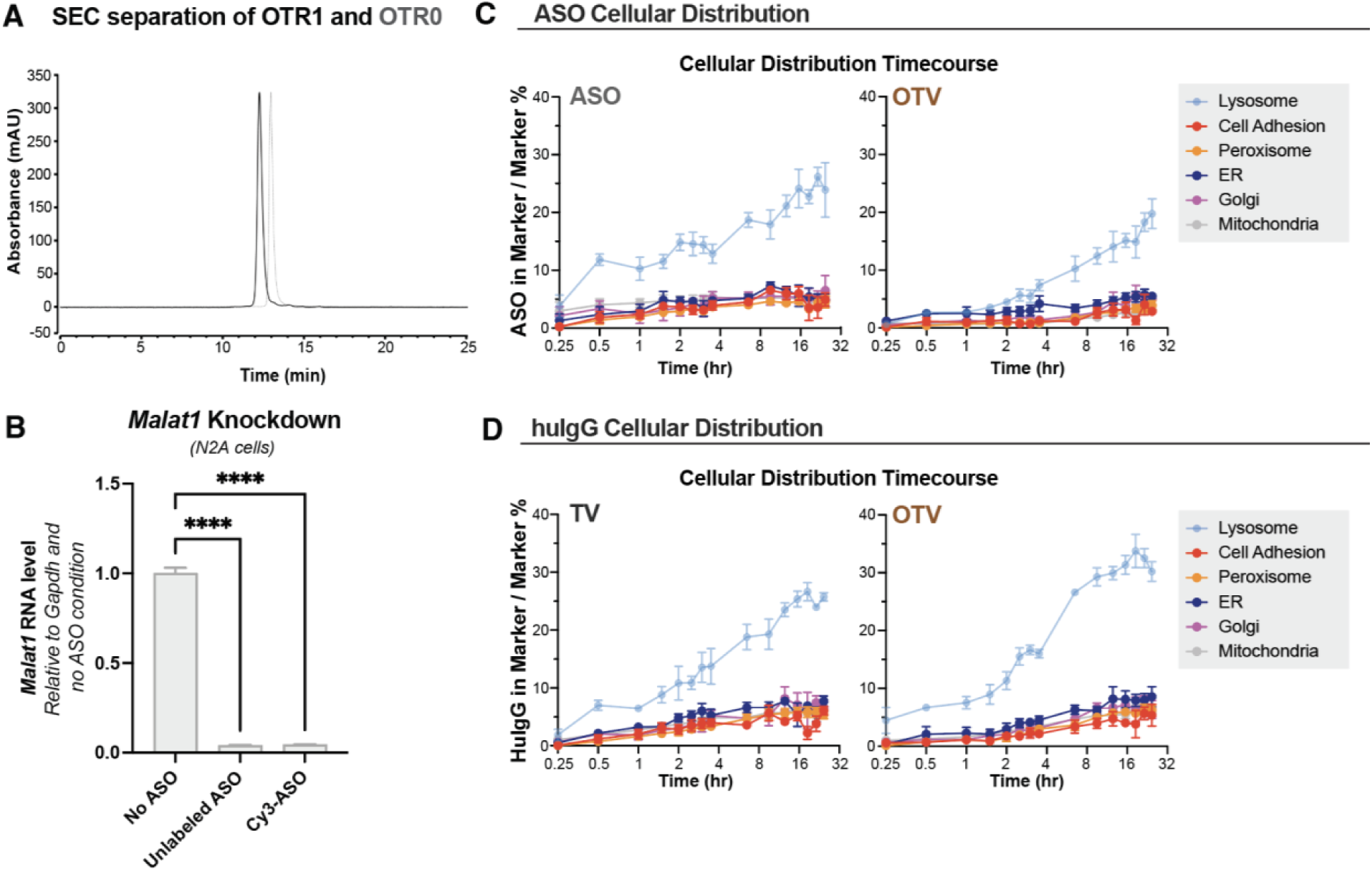
OTV purification, ASO efficacy, and time course of intracellular trafficking in SH-SY5Y cells. (**A**) SEC data showing separation between OTR1 (solid black line) and OTR0 (dashed gray line). **(B)** *Malat1* expression levels in cells treated with unlabeled ASO and Cy3-labeled ASO. **(C-D)** Live-cell accumulation over time of ASO (C) and huIgG (D) from naked ASO or OTV in indicated organelles by measuring Cy-3 (ASO) or Alexa-647 (huIgG) fluorescence intensity. Organelles were imaged using BacMam2.0 encoded GFP fusion marker proteins. Data are shown as means ± SEM; n = 3 independent experiments. One-way ANOVA adjusted for multiple comparisons using Sidak’s method for (B). ****p< 0.0001.

**Fig. S2.**
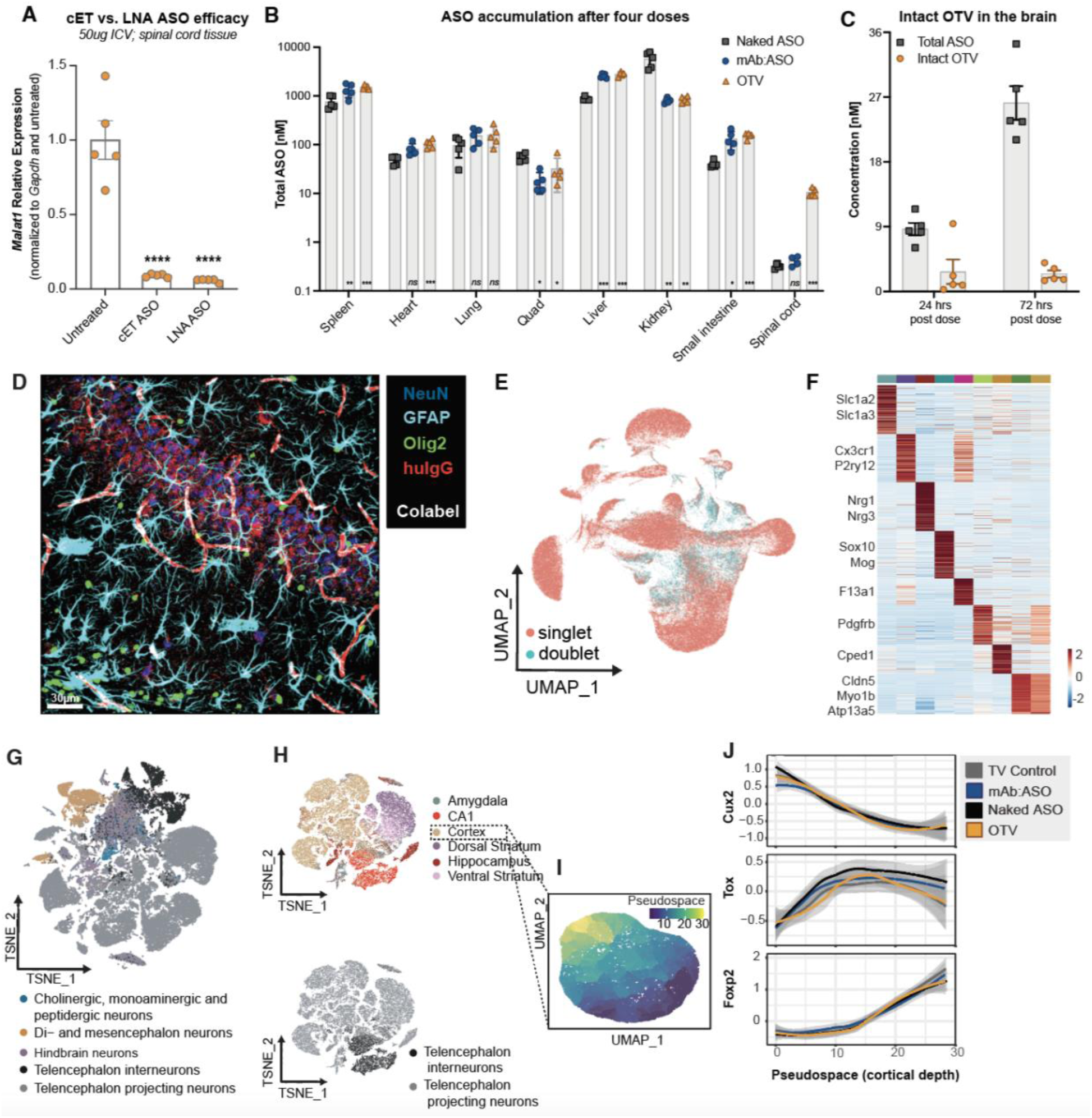
ASO efficacy, OTV pharmacokinetic response, and cortical snRNAseq pseudospace analysis. **(A)** *Malat1* knockdown in WT mouse spinal cord tissue after a 50 µg ICV injection of naked ASO containing either constrained-ethyl (cET) or LNA backbone modifications (Cohort 8). **(B)** ASO concentration in the TfR^mu/hu^ KI mouse spinal cord, quadricep muscle, liver, kidney, spleen, heart, lung, and intestine 72 hours after four doses of OTV (orange), naked ASO (grey) or mAb:ASO (blue). **(C)** Total ASO concentration (grey symbols) and intact OTV concentration (orange symbols) in the brain 24 hours after a single dose of OTV and 72 hours after four doses of OTV. **(D)** Image of the hippocampus after a single dose of OTV. Stained for huIgG (red), NeuN (blue), Gfap (turquoise), and Olig2 (green). Scale bar 30mm. (**E)** UMAP dimensionality reduction showing all nuclei identified as singlets (pink) or doublets (turquoise). (**F)** Heatmap showing average expression of 1800 marker genes across all predicted cell types excluding “glia”. (**G)** UMAP projection of all neurons, colored by predicted neuron subtype. (**H)** UMAP projections of all telencephalon neurons colored by predicted region (top) or broad subtype (bottom). **(I)** UMAP projection of cortical neurons generated using a subset of the expression matrix containing ∼500 genes with specific cortical layer expression patterns. Cells are colored according to pseudotime predicted using Monocle 3. **(J)** Local regression plots showing moving averages of scaled expression in each treatment group across pseudotime. Calculated with 500 randomly selected cells from each condition. Shaded regions show 95% confidence interval. Data are shown as means ± SEM. One-way ANOVA adjusted for multiple comparisons to the Untreated group using Dunnett’s method for (A). Data are shown as means ± SEM. Two-way ANOVA adjusted for multiple comparisons to the ASO group using Dunnett’s method for (B). ns p>0.05, ***p< 0.001 ****p< 0.0001.

**Fig. S3:**
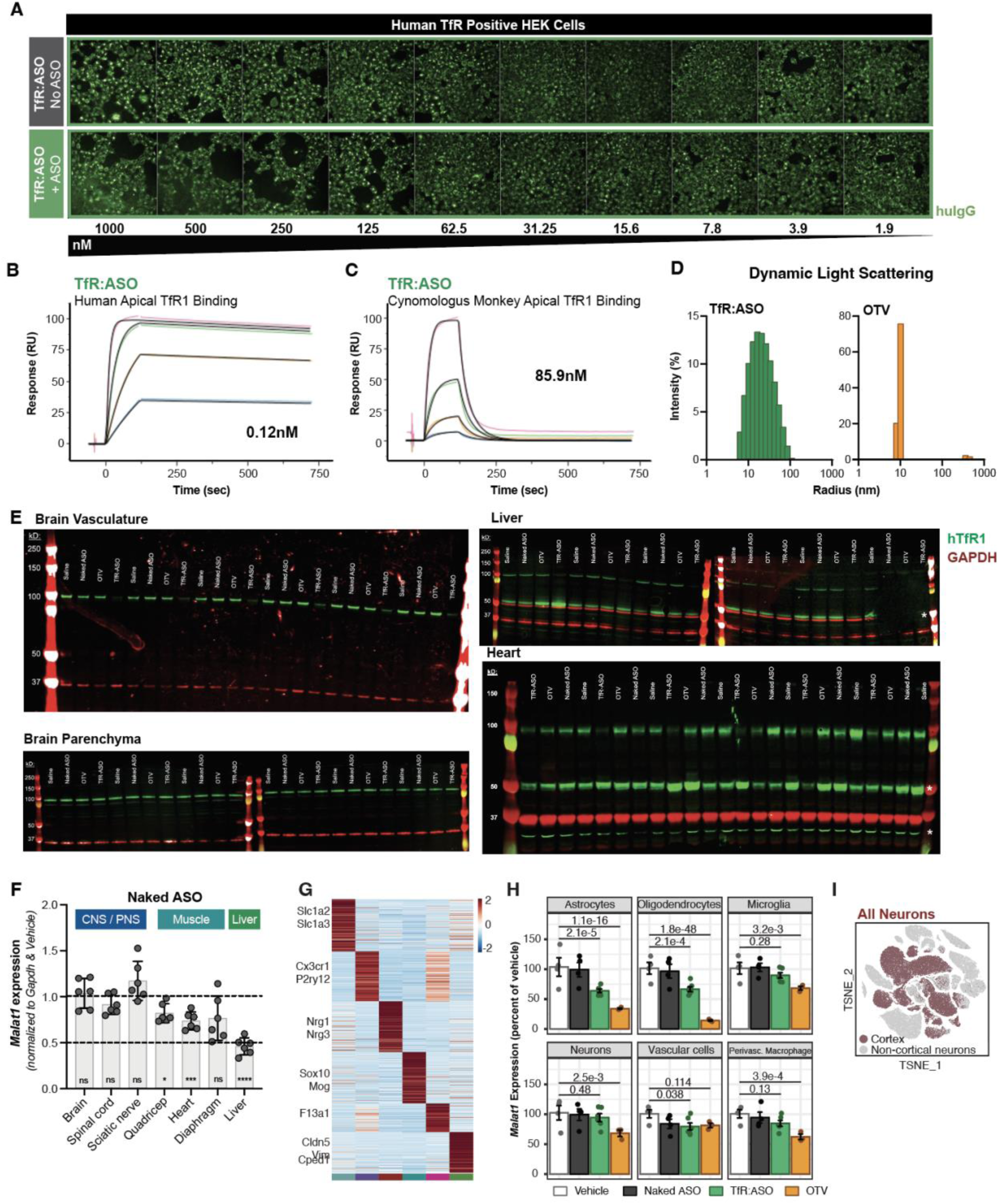
TfR:ASO molecule characterization and impact on hTfR1 protein and *Malat1* RNA levels. (**A)** No impact of ASO conjugation on high affinity, bivalent hTfR1 binding demonstrated by an *in vitro* uptake assay in HEK cells expressing hTfR1. **(B, C)** Binding of TfR:ASO to human (B) and cynomolgus monkey (C) TfR1 by SPR. (**D)** Dynamic light scattering of TfR:ASO (green) and OTV (orange) dosing solutions following incubation with full-length hTfR1 protein. (**E)** *Cohort 5:* Full western blot images showing hTfR1in green (band at ∼100 kDa) and Gapdh in red (band at ∼37 kDa). White asterisks indicate suspected nonspecific bands that do not correlate with any known form of hTfR and are not identified in either brain vasculature or parenchyma. (**F)** *Cohort 5: Malat1* knockdown in various tissues after treatment with naked ASO, normalized to *Gapdh* expression and displayed as a fold change relative to Saline-treated mice. (**G)** Heatmap showing average expression of 1200 marker genes across all predicted cell types excluding “glia”. (**H)** Boxplots showing pseudobulked scaled expression values for *Malat1* across cell types. Pseudobulk libraries were generated by summing counts for each cell type within mouse samples. Differential expression was performed in DESeq2 using Vehicle as the baseline comparison. P value numbers listed. **(I)** TSNE dimensionality reduction showing a subclustering of all telencephalon neurons. Neurons were identified as cortex or non-cortical through predictive label transfer of neuronal subtypes from the Mouse Brain Atlas. Data are shown as means ± SEM., Two-way ANOVA adjusted for multiple comparisons to the Saline group using Dunnett’s method for (F). ns p>0.05, * p<0.05, ***p< 0.001 ****p<0.0001

**Fig. S4.**
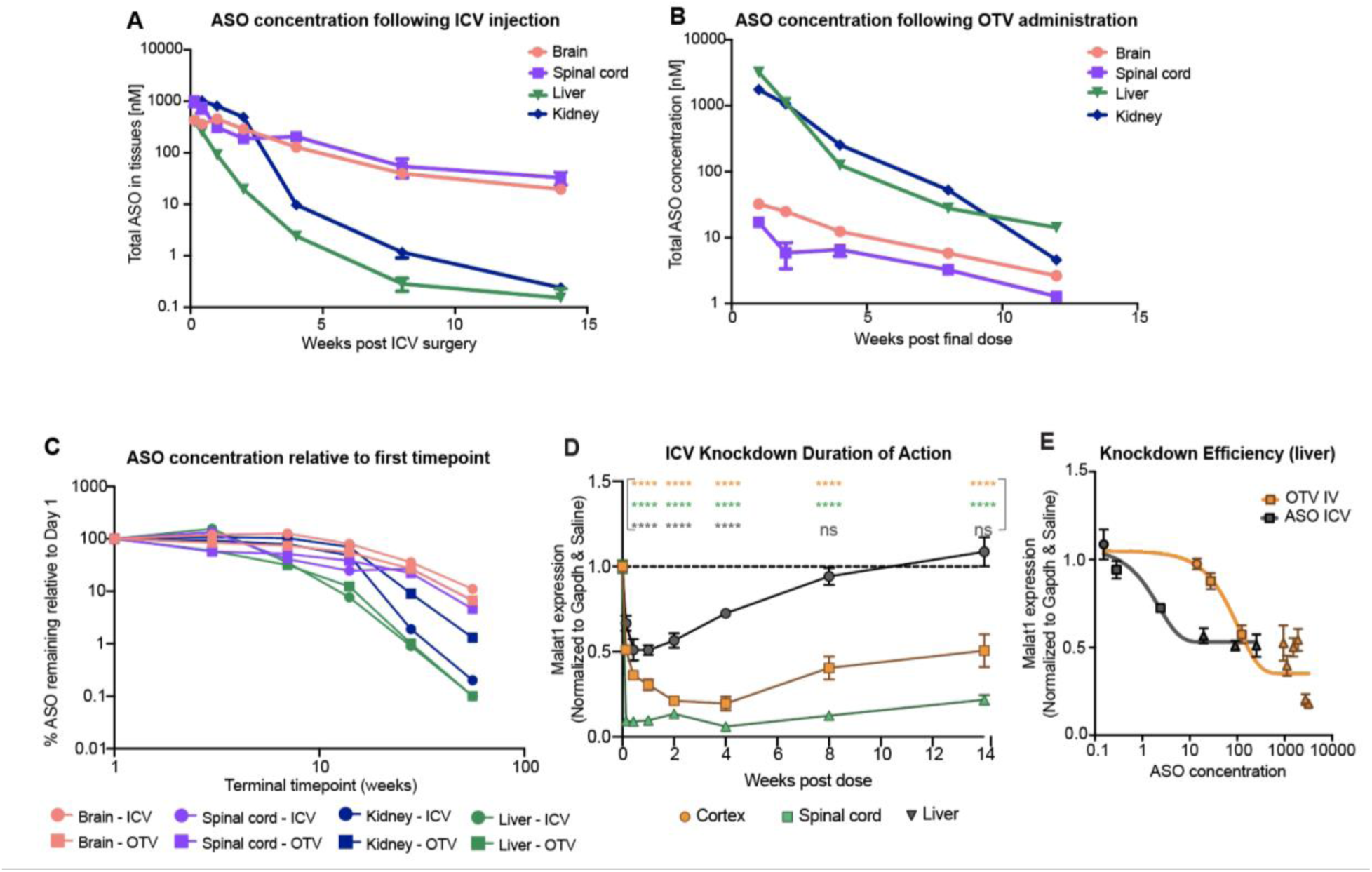
OTV preserves long ASO half-life and knockdown efficiency in the mouse brain. **(A-B)** ASO concentration in the brain (pink circle), spinal cord (purple square), liver (green triangle), and kidney (blue diamond) after ICV injection (A) or six IV doses (2X per week) of OTV (B). (**C)** ASO concentration in brain (pink), spinal cord (purple), liver (green), and kidney (blue) normalized to the first collection timepoint after ICV injection of naked ASO (circle) or six intravenous doses of OTV (square), demonstrating similar ASO half-life between OTV and ASO ICV conditions across tissues despite differences in starting concentration between the two conditions. (**D)** *Malat1* expression in the brain (orange), spinal cord (green), and liver (grey) at various timepoints after ICV injection. (**E)** Liver ASO concentration plotted against *Malat1* RNA expression as a measure of knockdown efficiency. Triangles represent collection timepoints under 2 weeks, squares represent 4- and 8-week collection timepoints, circles represent 12-and 14-week collection timepoints. Data are shown as means ± SEM. Two-way ANOVA adjusted for multiple comparisons to the Saline-dosed group using Dunnett’s method for (D). ns p>0.05, ***p< 0.001 ****p< 0.0001,

**Fig. S5:**
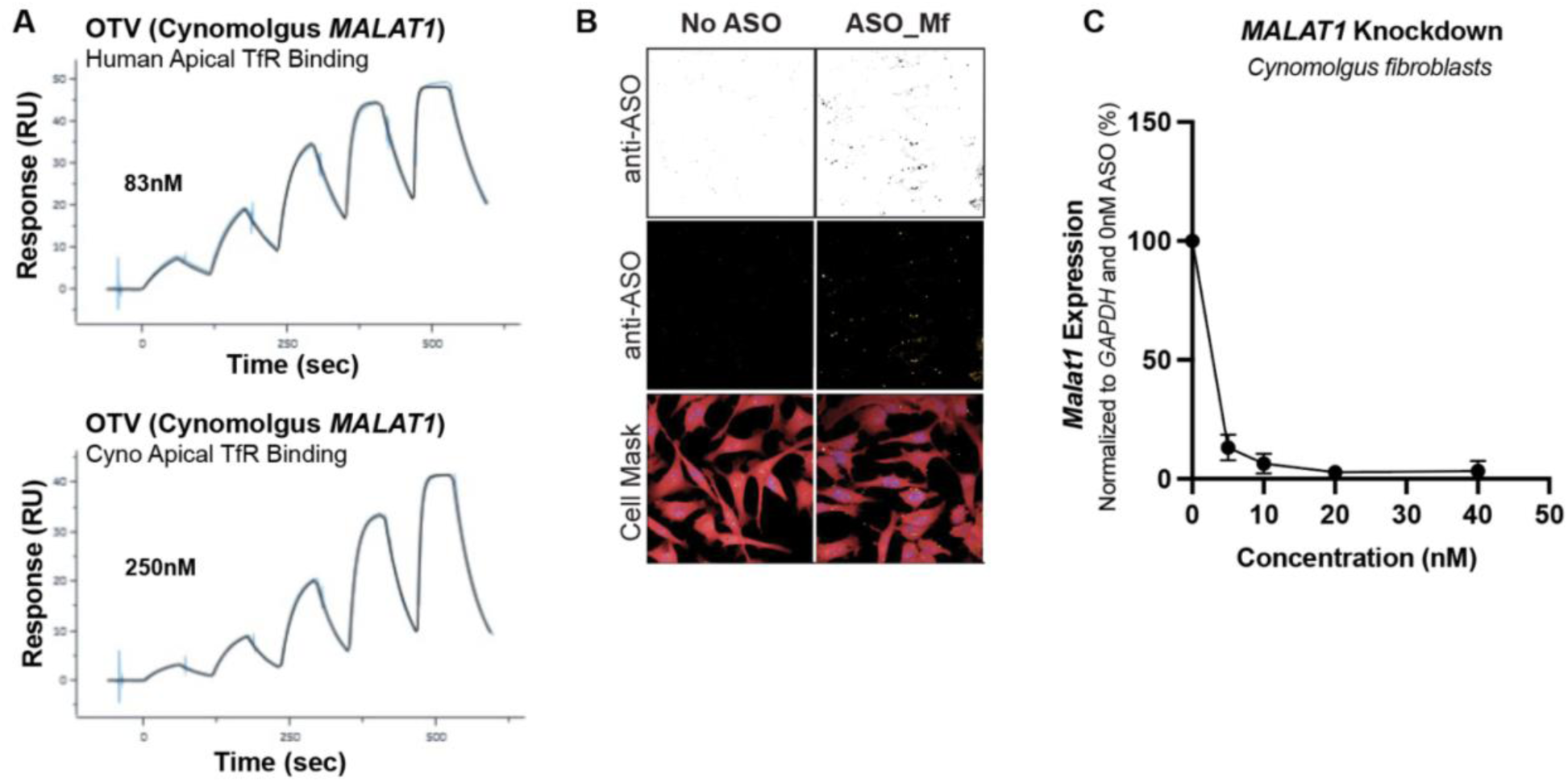
OTV compatible with cynomolgus monkey tissue is capable of uptake and robust knockdown *in vitro.* **(A)** Binding of OTV to human TfR1 (top) and cynomolgus monkey TfR1 (bottom) by surface plasmon resonance. **(B)** Uptake of 40nM full length *MALAT1* ASO^Mf^ into SHSY-5Y cells. Bottom panel shows the cell mask, middle panel is the raw fluorescence signal, and top panel is the inverted signal for visualization. (**C)** *MALAT1* RNA knockdown in cynomolgus monkey fibroblasts after incubation with ASO^Mf^. Data are shown as means ± SEM for three technical replicates.

**Fig. S6:**
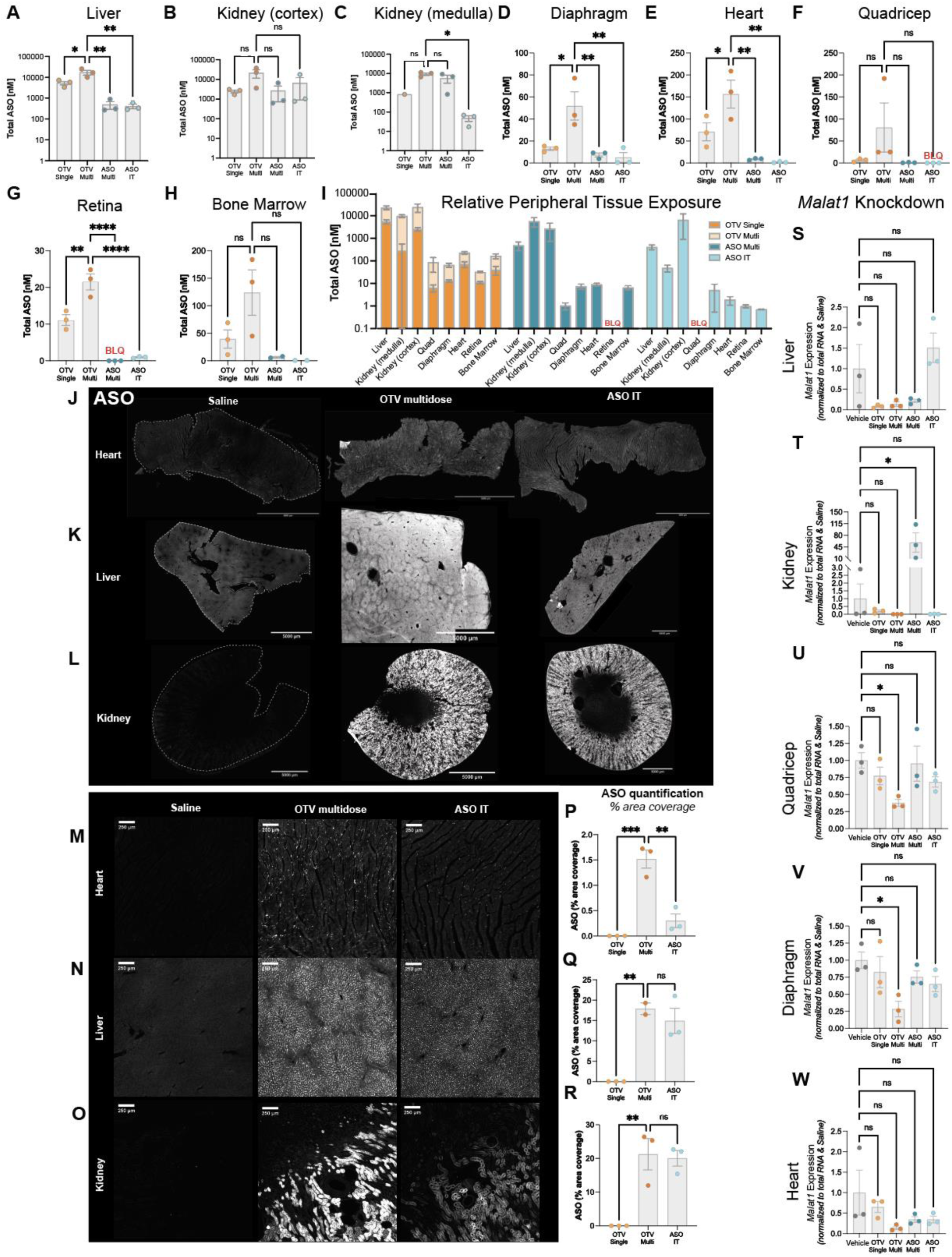
OTV results in significant ASO^Mf^ deposition and target knockdown in peripheral tissues in cynomolgus monkey. **(A-H)** Peripheral tissue ASO^Mf^ concentrations following a single IV dose of OTV (light orange), 4 doses of OTV (dark orange), 4 IV doses of naked ASO^Mf^ (dark blue) or a single IT dose of naked ASO^Mf^ (light blue). (**I)** Relative tissue concentrations of ASO^Mf^ across tissue and dosing paradigms. (**J-L)** Widefield images of ASO^Mf^ in heart, liver and kidney. (**M-O**) Higher magnification images of (J-L). (**P-R**) Quantification of (J-L). (**S-W)** Knockdown of *MALAT1* RNA in peripheral tissues. Data are shown as means ± SEM. One-way ANOVA adjusted for multiple comparisons to the Vehicle control group using Dunnett’s method. ns p>0.05, * p<0.05, ** p<0.01, *** p< 0.001 **** p< 0.0001

**Fig. S7:**
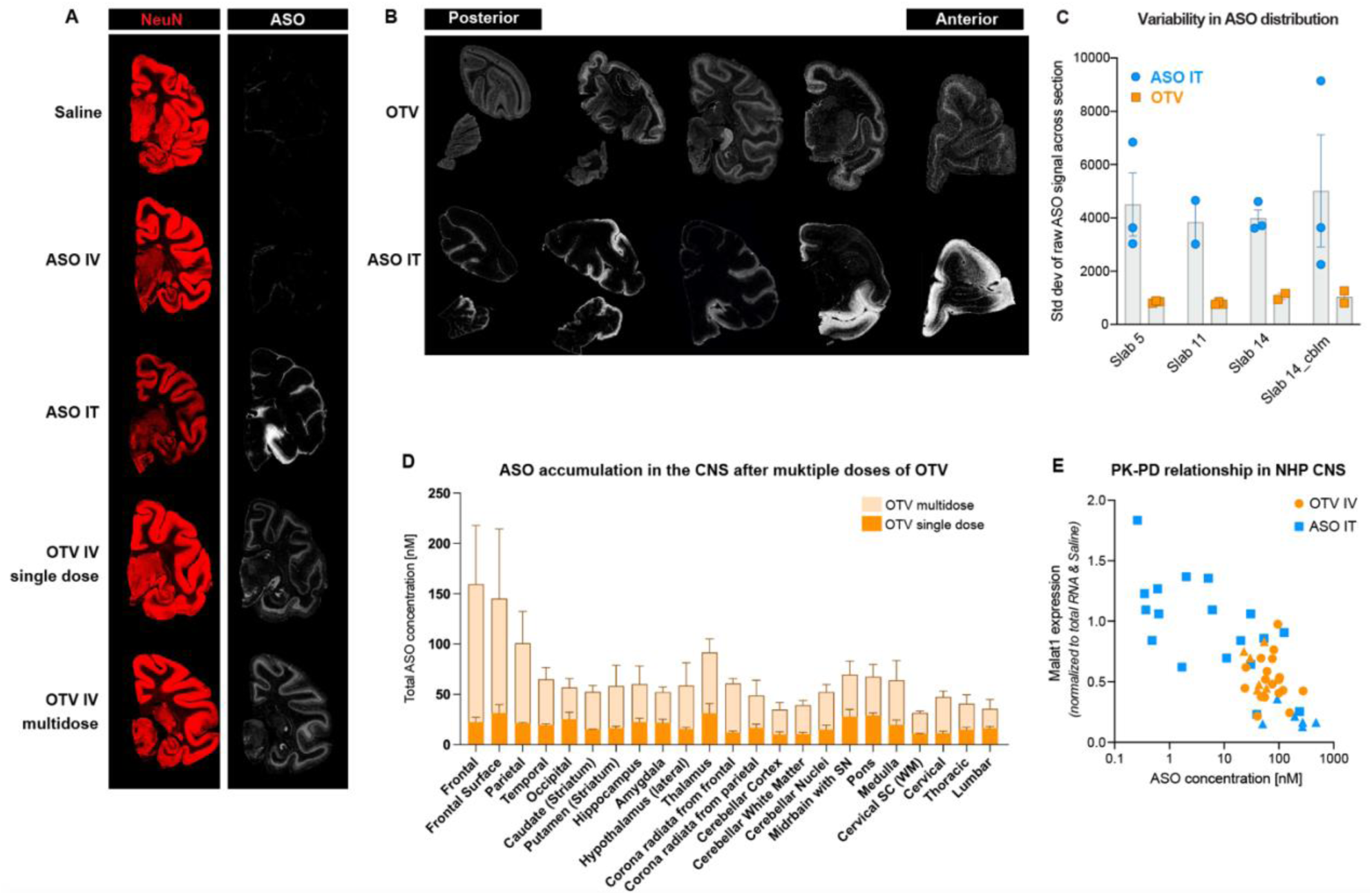
OTV provides cumulative and widespread ASO^Mf^ deposition in the cynomolgus monkey brain. **(A).** Representative images of immunofluorescent staining of NeuN (red) and ASO^Mf^ (white) in brain tissue from all experimental groups. (**B).** Representative images of immunofluorescent staining of ASO^Mf^ (white) across the AP (anterior – posterior) axis of the NHP brain. A subset of images are the same images shown in Figure 6. (**C).** Quantification of standard deviation in ASO^Mf^ signal across four slabs sampled throughout the brain. Lower standard deviation indicates greater homogeneity in ASO^Mf^ biodistribution. (**D).** ASO^Mf^ concentration in various regions of the CNS after a single (light orange) or four doses (dark orange) of OTV. n=3/group. For raw ASO concentration data for OTV single dose, OTV multi-dose, and IT groups, see Table S4. (**E**) Correlation between ASO concentration and *MALAT1* knockdown in the NHP CNS after IV OTV (orange) or ASO IT (blue) administration. Each point represents data from a single brain or spinal cord region from a single NHP. Triangles indicate spinal cord regions. Data are shown as means ± SEM.

**Fig. S8:**
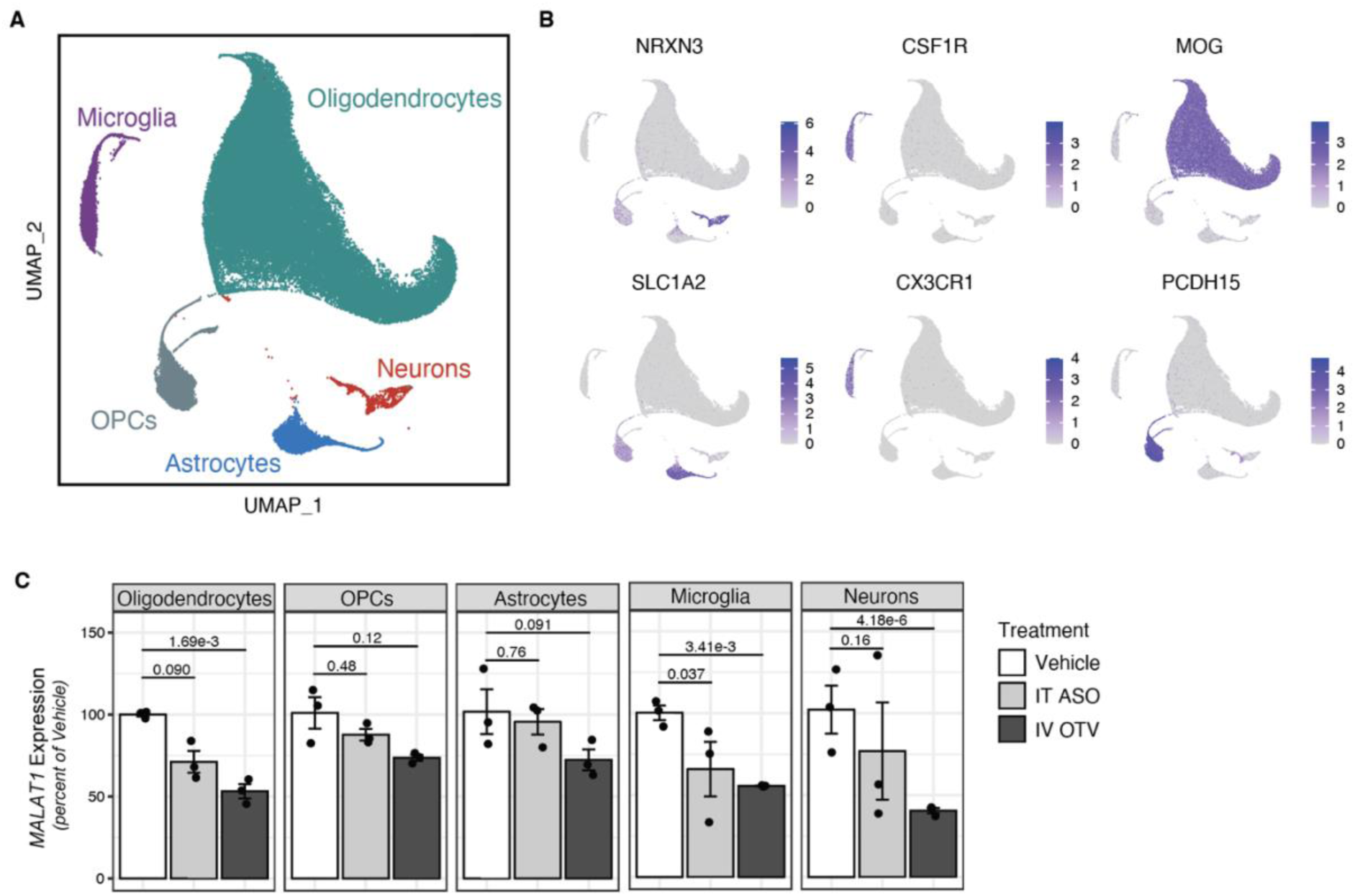
snRNAseq in cynomolgus monkey thalamic tissue. **(A)** UMAP dimensionality reduction showing all nuclei passing QC thresholds. (**B)** Cell type predictions were assigned via marker gene expression. (**C)** Boxplots showing pseudobulked scaled expression values for *MALAT1* across cell types. Pseudobulk libraries were generated by summing counts for each cell type within samples. Differential expression was performed in DESeq2 using Saline-treated as the baseline comparison. p-values listed.

**Table S1.** Pharmacokinetic profile of ASO^Mf^ and OTV dosed intravenously.

| Cohort | Analyte | C<sub>o</sub> (μmol/L) | C<sub>max</sub> (μmol/L) | AUC<sub>(0-inf)</sub> (hr*μmol/L) | CL (mL/day/kg) | t<sub>1/2</sub> (hr) |
| --- | --- | --- | --- | --- | --- | --- |
| ASO<sup>Mf</sup><br>IV | ASO<sup>Mf</sup> | NA | 0.83 (± 0.08) | 3.91 (± 0.55) | NA | NA |
| ASO<sup>Mf</sup><br>IT | ASO<sup>Mf</sup> | 3.76 (± 0.67) | NA | 0.99 (± 0.08) | 4.99E06 (± 4.08E05) | 7.36 (± 0.34) |
| OTV IV | ASO<sup>Mf</sup> | 4.61 (± 0.99) | NA | 91.06 (± 14.53) | 55.86 (± 8.30) | 21.07 (± 7.17) |
| OTV IV | huIgG | 5.15 (± 0.54) | NA | 216.98 (± 8.68) | 23.10 (± 0.90) | 45.69 (± 8.75) |

**Table S2.** Regions sampled throughout the NHP brain and spinal cord

|  |  |
| --- | --- |
| 1 | Frontal cortex |
| 2 | Frontal surface (top 2-3 mm of cortex) |
| 3 | Parietal cortex |
| 4 | Temporal/entorhinal cortex |
| 5 | Occipital cortex |
| 6 | Striatum containing caudate |
| 7 | Striatum containing putamen |
| 8 | Hippocampus |
| 9 | Amygdala |
| 10 | Hypothalamus (lateral) |
| 11 | Thalamus |
| 12 | Corona radiata (frontal) |
| 13 | Corona radiata (parietal) |
| 14 | Outer cerebellar cortex |
| 15 | Cerebellar white matter |
| 16 | Deep cerebellar nuclei |
| 17 | Midbrain with substantia nigra |
| 18 | Pons |
| 19 | Medulla |
| 20 | Cervical spinal cord white matter |
| 21 | Cervical spinal cord grey matter |
| 22 | Lumbar spinal cord grey matter |
| 23 | Thoracic spinal cord grey matter |

**Table S3.** ASO^Mf^ retained in CNS after OTV (IV) and ASO^Mf^ (IT) dosing.

|  | ASO<sup>Mf</sup> measured (nM) |  | Approx total tissue mass (g) | Total ASO<sup>Mf</sup> in tissue (mg) |  | Amount ASO<sup>Mf</sup> injected (mg) |  | % of injected ASO<sup>Mf</sup> retained |  |
| --- | --- | --- | --- | --- | --- | --- | --- | --- | --- |
|  | OTV (IV) | ASO<sup>Mf</sup> (IT) |  | OTV (IV) | ASO<sup>Mf</sup> (IT) | OTV (IV) | ASO<sup>Mf</sup> (IT) | OTV (IV) | ASO<sup>Mf</sup> (IT) |
| Brain<br>(avg ± SEM) | 70.50<br>(±6.38) | 40.95<br>(±7.67) | 70 | 0.26<br>(±0.02) | 0.15<br>(±0.03) | 9.70<br>(±0.38) | 4.00<br>(±0.00) | 2.72<br>(±0.50) | 3.82<br>(±1.94) |
| Spinal cord<br>(avg ± SEM) | 38.90<br>(±3.53) | 226.16<br>(±48.33) | 3.8 | 0.01<br>(±0.00) | 0.05<br>(±0.01) |  |  | 0.08<br>(±0.01) | 1.15<br>(±0.25) |
| ASO<sup>Mf</sup> Ratio in Brain: Spinal Cord | 1.81 | 0.18 | ASO<sup>Mf</sup> retained in CNS (mg) | 0.27 | 0.20 | ASO<sup>Mf</sup> retained in CNS (%) |  | 2.79 | 4.97 |

**Table S4.** Total ASO Concentration across Brain and Spinal Cord NHP Regions. Color gradient reflects high (red) to low (blue) total ASO concentration measured across brain regions.

| Total ASO (nM) |  |  |  |  |  |  |
| --- | --- | --- | --- | --- | --- | --- |
| Brain Region | IT Single Dose |  | OTV single dose |  | OTV multidose |  |
| Dose* | 4mg |  | 3-3.5mg |  | 9.2-10.6mg |  |
|  | Average | SEM | Average | SEM | Average | SEM |
| **Frontal** | 69.46 | 28.42 | 22.07 | 4.666 | 159.6 | 58.29 |
| **Frontal Surface** | 37.61 | 21.26 | 30.88 | 8.464 | 145.2 | 69.15 |
| **Parietal** | 99.07 | 69.55 | 20.94 | 0.5746 | 100.9 | 31.54 |
| **Temporal** | 36.39 | 24.87 | 19.03 | 1.183 | 65.11 | 11.58 |
| **Occipital** | 84.02 | 47.09 | 24.74 | 7.009 | 56.71 | 8.992 |
| **Caudate (Striatum)** | 2.27 | 1.901 | 14.58 | 0.302 | 52.64 | 6.174 |
| **Putamen (Striatum)** | 2.59 | 1.335 | 15.89 | 1.871 | 58.43 | 20.52 |
| **Hippocampus** | 41.11 | 28.4 | 22.15 | 3.641 | 60.27 | 17.83 |
| **Amygdala** | 65.94 | 36.46 | 21.05 | 3.703 | 52.21 | 5.162 |
| **Hypothalamus (lateral)** | 68.21 | 41.72 | 15.01 | 1.69 | 58.76 | 22.62 |
| **Thalamus** | 14.09 | 8.764 | 30.66 | 9.679 | 91.75 | 13.47 |
| **Corona radiata from frontal** | 3.417 | 3.098 | 11.51 | 1.517 | 60.83 | 4.788 |
| **Corona radiata from parietal** | 0.8033 | 0.4434 | 16.1 | 3.789 | 49.07 | 14.94 |
| **Cerebellar Cortex** | 65.13 | 52.1 | 9.741 | 2.588 | 34.85 | 7.16 |
| **Cerebellar White Matter** | 3.69 | 2.286 | 9.987 | 1.826 | 39.38 | 4.584 |
| **Cerebellar Nuclei** | 4.883 | 1.845 | 14.56 | 4.212 | 52.41 | 7.398 |
| **Midbrain with Substantia Nigra** | 74.18 | 57.44 | 27.33 | 7.146 | 69.71 | 13.39 |
| **Pons** | 17.01 | 12.78 | 28.43 | 2.551 | 67.61 | 12.2 |
| **Medulla** | 88.12 | 39.37 | 19.02 | 4.937 | 64.07 | 19.52 |
| **Spinal Cord Region** |  |  |  |  |  |  |
| **Cervical SC (White Matter)** | 121.9 | 38.42 | 10.32 | 0.5179 | 31.59 | 1.925 |
| **Cervical** | 112.7 | 42.85 | 10.75 | 2.139 | 47.54 | 5.794 |
| **Lumbar** | 331.2 | 140.2 | 14.47 | 2.399 | 35.64 | 9.391 |
| **Thoracic** | 338.8 | 68.51 | 15.7 | 1.904 | 40.84 | 8.831 |

\*Dose reflects exact amount of total ASO delivered either via direct CSF administration (IT) or intravenous (OTV) taking into account NHP body weight

**Table S5.** Detailed ASO synthesis parameters and protocol on 100 µM scale

| Process | Reagents | Parameters |
| --- | --- | --- |
| Solid support | UnyLinker CPG, 500 Å, 80 μmol/g | 100 μmol scale |
| Wash | Anhydrous acetonitrile (moisture content < 10 ppm) | 10 ml, 20 s |
| Detritylation | 3% Trichloroacetic acid (TCA) in DCM | 10 ml, 45 s |
| Coupling | 0.1 M phosphoramidites in ACN or ACN/DCM (1:1, v/v) | 2.5 ml amidite,<br>5 ml ETT, 6 min |
|  | 0.25 M ETT in acetonitrile |  |
| Oxidation | 0.02 M I<sub>2</sub> in THF/Pyridine/H<sub>2</sub>O (7:2:1) | 10 ml, 2 min |
| Sulfurization | 0.2 M PADS in 50% pyridine/50% acetonitrile | 10 ml, 4 min |
| Capping | Cap Mix A: THF/Pyridine/Ac<sub>2</sub>O (8:1:1) | 2.5 ml Cap A, 2.5 ml<br>Cap B, 40 s |
|  | Cap Mix B: 16% Methyl imidazole in THF |  |
| Step | Process |  |
| Initialization | a) Wash 3×<br>b) Detritylation 3× |  |
| Elongation | 1<sup>st</sup> monomer | a) Detritylation 4×<br>b) Wash 3×<br>c) Coupling 4×<br>d) Oxidation (or sulfurization)<br>e) Capping<br>f) Wash 3× |
|  | 2<sup>nd</sup> to end monomer | a) Detritylation 4×<br>b) Wash 3×<br>c) Coupling 2×<br>d) Oxidation (or sulfurization)<br>e) Capping<br>f) Wash 3× |
| Finalization | Wash 3x |  |

**Table S6.** Prep-HPLC purification conditions

|  |  |
| --- | --- |
| Instrument | GILSON-281/322/159 |
| Signal | 260 nm |
| Column | Waters Xbridge BEH C18 19*150 mm |
| Mobile phase A | 240 mM HFIP and 7 mM TEA in 95% water/5% MeOH |
| Mobile phase B | 240 mM HFIP and 7 mM TEA in MeOH |
| Flow rate | 15 ml/min |
| Column temperature | 50 °C |
| Gradient | 0–3 min (10% B), 3–25 min (10% - 30%), 100% B hold time: 2 min |
| Amount per injection | 100 mg crude ASO |

**Table S7:** Detailed Mouse Information.

| **Cohort 1 (Fig. 3A)** |  |  |  |  |  |  |  |
| --- | --- | --- | --- | --- | --- | --- | --- |
| **Treatment** | **Genotype** | **Sex (#)** | **Age\*** | **Dose\*\*** | **Route** | **Single/Multi-Dose** | **Tissue Timepoint** |
| OTV | WT | F (3) | 2.7 | 25 | IV | Single | NA |
| Naked ASO | WT | F (3) | 2.7 | 0.9 | IV | Single | NA |
| **Cohort 2 (Fig. 3C; fig. S2C-D)** |  |  |  |  |  |  |  |
| **Treatment** | **Genotype** | **Sex (#)** | **Age\*** | **Dose\*\*** | **Route** | **Single/Multi-Dose** | **Tissue Timepoint** |
| TV Control | TfR<sup>mu/hu</sup> KI | F (2), M (3) | 2.3-2.6 | 58 | IV | Single | 24hr |
| mAb:ASO | TfR<sup>mu/hu</sup> KI | F (3), M (2) | 2.3-2.6 | 60 | IV | Single | 24hr |
| Naked ASO | TfR<sup>mu/hu</sup> KI | F (3), M (2) | 2.3-2.6 | 2.4 | IV | Single | 24hr |
| OTV | TfR<sup>mu/hu</sup> KI | F (2), M (3) | 2.3-2.6 | 60 | IV | Single | 24hr |
| **Cohort 3 (Fig. 3C-H; fig. S2B, E-J)** |  |  |  |  |  |  |  |
| **Treatment** | **Genotype** | **Sex (#)** | **Age\*** | **Dose\*\*** | **Route** | **Single/Multi-Dose** | **Tissue Timepoint** |
| TV Control | TfR<sup>mu/hu</sup> KI | F (3), M (2) | 2.3-2.6 | 58 | IV | Multi (4x) - weekly | 72hr post last dose |
| mAb:ASO | TfR<sup>mu/hu</sup> KI | F (3), M (2) | 2.3-2.6 | 60 | IV | Multi (4x) - weekly | 72hr post last dose |
| Naked ASO | TfR<sup>mu/hu</sup> KI | F (3), M (2) | 2.3-2.6 | 2.4 | IV | Multi (4x) - weekly | 72hr post last dose |
| OTV | TfR<sup>mu/hu</sup> KI | F (3), M (2) | 2.3-2.6 | 60 | IV | Multi (4x) - weekly | 72hr post last dose |
| **Cohort 4 (Fig. 4E-H)** |  |  |  |  |  |  |  |
| **Treatment** | **Genotype** | **Sex (#)** | **Age\*** | **Dose\*\*** | **Route** | **Single/Multi-Dose** | **Tissue Timepoint** |
| Saline | TfR<sup>mu/hu</sup> KI | F (4) | 2.3 | NA | IV | Single | 24hr |
| Naked ASO | TfR<sup>mu/hu</sup> KI | F (4) | 2.3 | 2 | IV | Single | 24hr |
| TfR:ASO | TfR<sup>mu/hu</sup> KI | F (4) | 2.3 | 50 | IV | Single | 24hr |
| OTV | TfR<sup>mu/hu</sup> KI | F (4) | 2.3 | 50 | IV | Single | 24hr |
| **Cohort 5 (Fig. 4C-D, I-N; fig. S3E-I)** |  |  |  |  |  |  |  |
| **Treatment** | **Genotype** | **Sex (#)** | **Age\*** | **Dose\*\*** | **Route** | **Single/Multi-Dose** | **Tissue Timepoint** |
| Saline | TfR<sup>mu/hu</sup> KI | F (6) | 3 | NA | IV | Multi (4x) - weekly | 72hr post dose |
| Naked ASO | TfR<sup>mu/hu</sup> KI | F (6) | 3 | 2 | IV | Multi (4x) - weekly | 72hr post dose |
| TfR:ASO | TfR<sup>mu/hu</sup> KI | F (6) | 3 | 50 | IV | Multi (4x) - weekly | 72hr post dose |
| OTV | TfR<sup>mu/hu</sup> KI | F (5) | 3 | 50 | IV | Multi (4x) - weekly | 72hr post dose |
| **Cohort 6 (Fig. 5A-C,E,F; fig. S4B,C,E)** |  |  |  |  |  |  |  |
| **Treatment** | **Genotype** | **Sex (#)** | **Age\*** | **Dose\*\*** | **Route** | **Single/Multi-Dose** | **Tissue Timepoint** |
| Saline | TfR<sup>mu/hu</sup> KI | F (3) per timepoint | 3 | NA | IV | Multi (6x) - 2x wk | 1wk, 2wk, 4wk, 8wk, 12wk |
| OTV | TfR<sup>mu/hu</sup> KI | F (5) | 3 | 25 | IV | Multi (6x) - 2x wk | 1 wk post dose |
| OTV | TfR<sup>mu/hu</sup> KI | F (5) | 3 | 25 | IV | Multi (6x) - 2x wk | 2 wks post dose |
| OTV | TfR<sup>mu/hu</sup> KI | F (5) | 3 | 25 | IV | Multi (6x) - 2x wk | 4 wks post dose |
| OTV | TfR<sup>mu/hu</sup> KI | F (5) | 3 | 25 | IV | Multi (6x) - 2x wk | 8 wks post dose |
| OTV | TfR<sup>mu/hu</sup> KI | F (5) | 3 | 25 | IV | Multi (6x) - 2x wk | 12 wks post dose |
| **Cohort 7 (Fig. 5B,D,F; fig. S4A,C-E)** |  |  |  |  |  |  |  |
| **Treatment** | **Genotype** | **Sex (#)** | **Age\*** | **Dose\*\*** | **Route** | **Single/Multi-Dose** | **Tissue Timepoint** |
| Saline | WT | F (3) per timepoint | 2.6 | NA | ICV | Single | 24hr, 72hr, 1wk, 2wk, 4wk, 8wk, 14wk |
| Naked ASO | WT | F (5) | 2.6 | 50ug | ICV | Single | 24hr |
| Naked ASO | WT | F (5) | 2.6 | 50ug | ICV | Single | 72hr |
| Naked ASO | WT | F (4) | 2.6 | 50ug | ICV | Single | 1week |
| Naked ASO | WT | F (5) | 2.6 | 50ug | ICV | Single | 2 week |
| Naked ASO | WT | F (5) | 2.6 | 50ug | ICV | Single | 4 week |
| Naked ASO | WT | F (5) | 2.6 | 50ug | ICV | Single | 8week |
| Naked ASO | WT | F (5) | 2.6 | 50ug | ICV | Single | 14week |
| **Cohort 8 (fig. S2A)** |  |  |  |  |  |  |  |
| **Treatment** | **Genotype** | **Sex (#)** | **Age\*** | **Dose\*\*** | **Route** | **Single/Multi-Dose** | **Tissue Timepoint** |
| Saline | WT | F (5) | 6-7 | NA | ICV | Single | 72hr |
| LNA ASO | WT | F (5) | 6-7 | 50ug | ICV | Single | 72hr |
| cET ASO | WT | F (5) | 6-7 | 50ug | ICV | Single | 72hr |

\*Age is reported in months at the time of first dose
\*\*Dose is in mg/kg for IV; indicates dose of the whole molecule, not just the ASO portion - doses were selected for ASO molar equivalency

**Table S8:** Detailed NHP Information.

| NHP Cohort (Fig. 6; fig. S6-8) |  |  |  |  |  |  |  |
| --- | --- | --- | --- | --- | --- | --- | --- |
| Treatment | Sex (#) | Weight (kg) | Age (yrs) | Dose | Route | Single or Multi-Dose | Tissue Timepoint |
| ASO<sup>Mf</sup> | F (3) | 3.0-3.6 | 3 | 4mg | IT | Single | 2 weeks |
| OTV | F (3) | 2.3-3.2 | 2.7-3.7 | 30mg/kg | IV | Single | 1 week |
| Saline | F (3) | 2.0-2.4 | 2.5-2.6 | NA | IV | Multi-Dose (4x) - 2x a week | 2 weeks post last dose |
| OTV | F (3) | 2.1-2.4 | 2.6-2.7 | 30mg/kg | IV | Multi-Dose (4x) - 2x a week | 2 weeks post last dose |
| ASO<sup>Mf</sup> | F (3) | 2.9-3.6 | 3 | 1.1mg/kg | IV | Multi-Dose (4x) - 2x a week | 2 weeks post last dose |

